# Biological volume-EM with a focused Ga^+^ beam depends on forming radiation- resistant Ga-rich layer at block face

**DOI:** 10.1101/2024.09.16.613321

**Authors:** Maria A. Aronova, Zijie Yang, Joshua C. Kim, Cameron M. Baenen, Samuel J. Fulton, Guofeng Zhang, Xiaobing Chen, Richard D. Leapman

**Author notes:** Corresponding Author Mail: Richard D. Leapman, Ph.D. NIBIB, NIH Building 13, Room 3N17 13 South Drive Bethesda MD 20892. Co-first authors (equal contributions) (Maria A. Aronova, Zijie Yang, Richard D. Leapman).

## Abstract

Volume electron microscopy (vEM) enables biologists to visualize nanoscale 3D ultrastructure of entire eukaryotic cells and tissues processed by heavy atom staining and plastic embedding. The vEM technique with the highest resolution is focused ion-beam scanning electron microscopy (FIB-SEM), which provides nearly isotropic (∼5-10 nm) spatial resolution with a 1.5 keV beam at fluences up to 10,000 e^-^/nm^2^. However, it is still not understood how such resolution is achievable, because serial block-face (SBF)-SEM, which incorporates an in-situ ultramicrotome instead of a Ga^+^ FIB beam, results in radiation-induced collapse of similar specimen blocks at fluences of only ∼20 e^-^/nm^2^. In this study we demonstrate that FIB-SEM implants a thin surface layer of concentrated Ga^+^ ions, which significantly mitigates electron beam-damage, restricts the depth from which backscattered electrons are detected, and prevents specimen charging and collapse. Furthermore, we show that the depth resolution in *z*, perpendicular to block-face in FIB-SEM, is substantially higher than predicted by Monte Carlo modeling of the backscattered signal when Ga implantation is not included.

## Introduction

The advent of techniques for 3D volume electron microscopy (vEM) over the past two decades has transformed ultrastructural imaging of eukaryotic cells and tissues^1–4^. These methods require that the specimen be fixed, stained with heavy elements, and embedded in polymerized plastic. Ultrastructure can then be imaged in 3D from volumes ranging from 10^4^ µm^3^ encompassing single eukaryotic cells, to 10^9^ µm^3^ containing complex tissues. The journal *Nature* featured vEM as one of its seven technologies to watch in 2023, as tools and techniques that are poised to have an outsized impact on science in the coming year^5^. The technique having highest resolution, and suitable for imaging entire eukaryotic cells and tissues at a near isotropic resolution of ∼10 nm, is focused ion beam scanning electron microscopy (FIB-SEM), in which a Ga^+^ ion beam ablates very thin (3-10 nm) layers from a specimen block face, while a backscattered electron image is collected^6,7^. This process is repeated to acquire desired volumetric data. Such data sets reveal heavy atom-stained ultrastructure in 3D at a resolution of 5-10 nm. However, the reason why FIB- SEM is indeed capable of providing such high-resolution cellular structure has not yet been fully explained. Specifically, several aspects of the data collection in the FIB-SEM have appeared inconsistent with what is known about interactions between the electron probe and plastic- embedded stained biological specimens. Specifically, why can the sample blocks withstand such high electron fluences greater than 10^4^ e/nm^2^ without physical collapse^8^? Why is the structural information contained within the surface layer of 5-10 nm for 1.5 keV primary electrons^9^? And why doesn’t visible surface charge build-up occur^10^? Here we have performed experiments as well as simulations of ion and electron scattering that answer these questions through the discovery of an implantation layer containing a high concentration of Ga^+^ within ∼25 nm of the block’s surface.

To put these apparent inconsistencies in a wider context, it is helpful to consider briefly the various vEM techniques that are now beginning to find wide use. In the automated tape-collecting ultramicrotome (ATUM) thin sections are cut in the form of ribbons that are transferred first to tape and then to silicon wafers^4^; these arrays of sections are imaged in the scanning electron microscope (SEM) using a backscattered electron detector that is sensitive to local concentrations of heavy-atom stain (Os, Pb, and U), which provides contrast from the 3D cellular ultrastructure, equivalent to that obtained by conventional transmission electron microscopy (TEM). Although the ATUM enables the acquisition of very large ultrastructural volumes, e.g., cubic millimeters of brain tissue, with a lateral (*x*, *y*) spatial resolution of ∼5 nm, the spatial resolution in the *z*-direction (perpendicular to the block-face) is limited to ∼30 nm, corresponding to the minimum slice thickness that can be cut by the diamond knife of the ATUM’s ultramicrotome.^4^

An alternative to the ATUM is serial block-face SEM (SBF-SEM)^11–16^, in which the ultramicrotome is incorporated into the specimen stage of the SEM, enabling thin slices to be removed successively from the block-face in situ, in the vacuum of the SEM. In SBF-SEM the thin sectioned slices are discarded, and the fresh block-face is imaged using the backscattered signal generated by the electron beam whose energy is typically between 1.2 and 1.5 keV. The SBF-SEM has the advantage of avoiding distortions of the sections that can occur in the ATUM, but the *z*-resolution is again limited by the minimum slice thickness of ∼30 nm that can be removed from the block. It is also found that the block-face in the SBF-SEM is prone to electrical charging that distorts the images, although this can now be mitigated by use of a focused charge compensation device that neutralizes the charge build-up. A bigger limitation of the SBF-SEM is that the maximum electron fluence is only ∼20 electrons per nm^2^, beyond which the surface of the block undergoes radiation damage resulting in collapse. This limits the signal-to-noise ratio for detecting ultrastructural features in the image, even though the contrast of those features is high.

In addition, electron scattering simulations show that an incident probe of 1.5 keV generates a backscattered electron signal originating from a depth of at ∼25 nm into the block, suggesting that the *z*-resolution for the FIB-SEM should be ∼25 nm. In the Zeiss Crossbeam 550 FIB-SEM used in the current study, it is possible in principle to exclude electrons originating from a deeper layer in the specimen by filtering the energy of backscattered electrons using an energy-selective backscattered (ESB) detector^17^. This is facilitated by applying a bias voltage to an electrically isolated grid situated in front of the detector to repel electrons emerging from the specimen after having lost energy through multiple inelastic scattering deeper inside the specimen block. Alternatively, a bias voltage can be applied to the specimen itself so that electrons undergoing energy loss are excluded from entering the backscattered electron detector^18^. In practice, we have found that energy-selection alone is insufficient to provide spatial resolution of ∼5 to 10 nm in the *z*-direction comparable to the attainable resolution in the x-y plane, as we describe later. Therefore, we believe that achievement of isotropic resolution is not predicted in the absence of Ga implantation.

Acquiring 3D ultrastructural images from plastic-embedded stained biological structures using FIB-SEM requires several in-situ preparation steps that are performed under vacuum in the specimen stage. To understand the interaction of the Ga^+^ and e^-^ beams with the specimen, it is useful to consider the specimen stage set-up (Fig. 1a). Firstly, it is necessary to create a block-face in the *x-y* plane of the scanned focused ion beam, and to ensure that this block-face is accessible to the scanning electron beam. In the Zeiss Crossbeam 550 FIB-SEM used for this study, the FIB beam is at an angle of 54° to the electron beam. The specimen block’s surface is therefore tilted to an angle of 54° towards the FIB beam, and a block-face (in the *x-y* plane) perpendicular to the block’s surface (*x-z* plane) is prepared for imaging. Secondly, creation of the block-face requires deposition of a protective pad of a heavy element (platinum in our instrument) using the FIB beam to decompose an evaporated organometallic Pt compound. The protective pad avoids non-uniform ion beam milling, which is also referred to as ‘curtaining’^19^. The pad also allows the fabrication of registration marks by removing thin diagonally oriented grooves in the plane of the deposited Pt, which are then filled with carbon from an organic liquid released into the vacuum of the FIB- SEM. This yields slice-by-slice calibration markers that enable direct measurement of distance perpendicular to the block-face during the FIB-SEM volume acquisition, thus providing the thickness of each slice as well as autofocus and autostigmations together with lateral tracking in *x-y*. Finally, we can expose the block-face by milling a trench in the specimen block, allowing access to the electron beam as illustrated in Fig. 1a,b. The trench material is removed rapidly by increasing the FIB current to ∼30 nA, which forms a triangular profile in the *y-z* plane (Fig. 1b) caused by ejection of plastic-embedding material along with Ga^+^ ions. After the trench is formed, the block-face is polished by reducing the ion current to ∼700 pA, and thin layers are successively ablated from the volume of interest.

**Figure 1.**
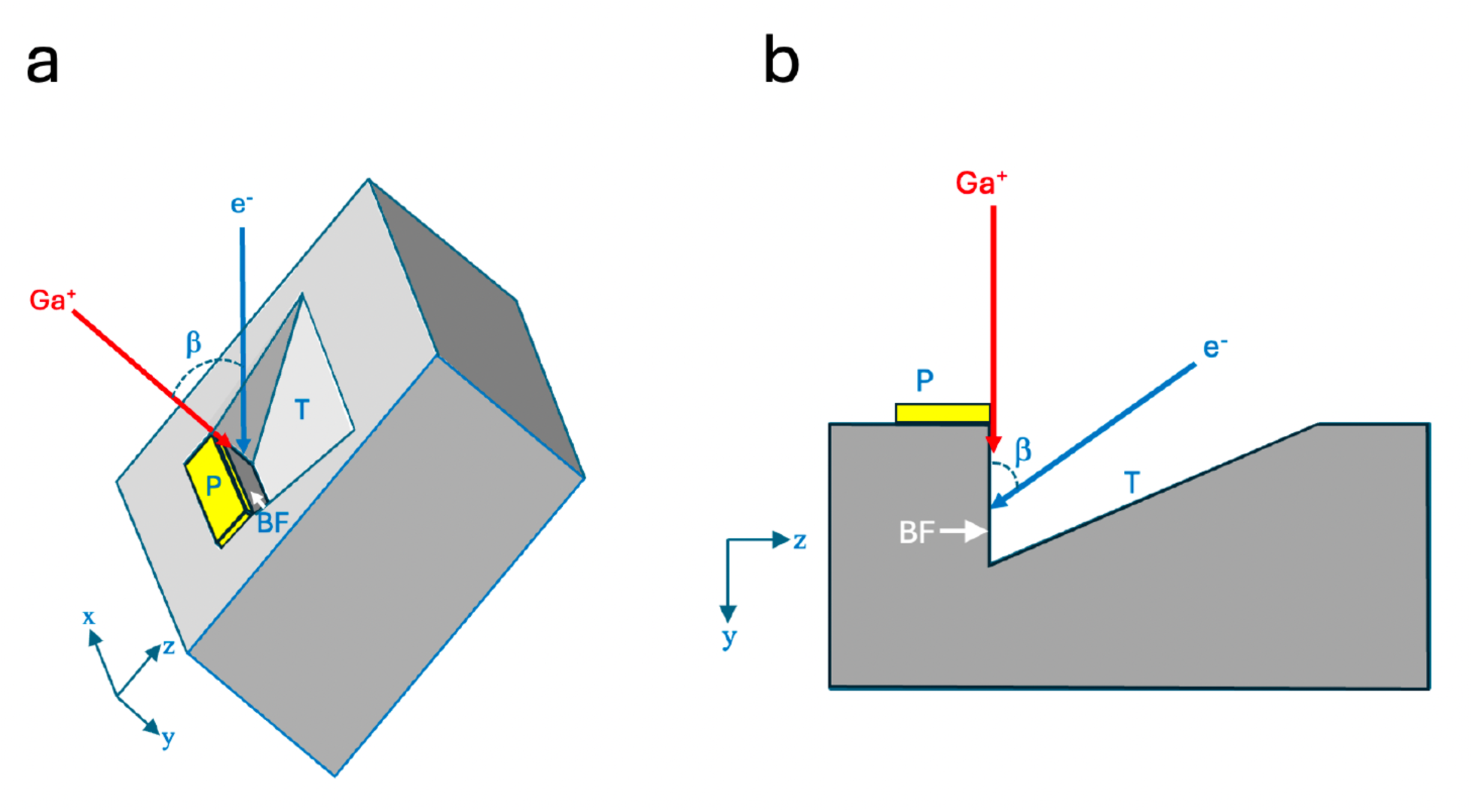
Principle of FIB-SEM, as applied to fixed, stained, and embedded biological specimen block. **(a,b)** Schematic illustrations showing, respectively, the *in-situ* preparation geometry in three dimensions (lateral *x*, height *y*, and depth *z*), and in two dimensions (as viewed along the *x*-axis in the *y*-*z* plane). The specimen block is tilted to angle β, enabling an incident 30- keV Ga^+^ beam at angle β from the vertical axis of the electron column to mill a trench (T), which allows geometrical access to the incident 1.5-keV electron probe (e^-^) as well as collection of backscattered electrons by an energy-selected backscattered (ESB) detector (**a**). A micrometer- thick platinum pad (P) is deposited on top of the volume to be imaged by scanning the focused ion beam while organometallic Pt is released into the vacuum of the SEM in proximity to the specimen block. Diagonal line patterns are then etched in the Pt pad by the ion beam and filled in with a layer of deposited carbon, which provides an internal calibration of the milling depth along the *z*- axis (not shown). The Pt pad prevents irregular cutting of the block-face (BF) by the ion beam. Although the trench is milled normal to block surface, material released in the direction opposite to the uncut block forms a downward sloping surface ending at the foot of the block face (**b**). The milled trench is trapezoidal, it is narrower at the block face than at the start of the trench furthest from the block face, to facilitate collection of the backscattered electrons (BSEs). After the Pt pad is deposited and the trench has been prepared, the volume data is acquired by continuous, simultaneous operation of the crossed Ga^+^ and e^-^ beams.

We illustrate the 3D ultrastructure obtained from FIB-SEM with a sample of cultured dissociated rat hippocampal neurons that is fixed and stained before embedding in Epon/Araldite (Fig. 2). Examination of neurite plasma membranes, endoplasmic reticulum membranes, and synaptic vesicle membranes in Fig. 2a, 2b, 2c, and 2d indicates widths of 5-10 nm in the *x*-*y* plane, with a slightly lower resolution of ∼10 nm in the *y*-*z* plane, which we attribute to penetration of the 1.5 keV electron probe ∼10 nm into the block. The 3D ultrastructural images in the FIB-SEM clearly enable visualization of circular cross sections through synaptic vesicles in the *x*-*z* or *y*-*z* plane (Fig. 2e) from which surface-rendered models can be extracted. These individual synaptic vesicles can be tracked throughout the volume, as illustrated by the presynaptic terminal containing 24 *z*-slices spaced 3 nm apart in Fig. 2f; a vesicle of diameter ∼50 nm indicated by a yellow arrow appears in the 9-nm slice and persists until the 66-nm slice. As we shall see in Fig. 8, the sharpness of the vesicle membrane in the z-direction (Fig. 2e and 2f) cannot be explained by backscattered electron simulations in the absence of Ga implantation.

**Figure 2.**
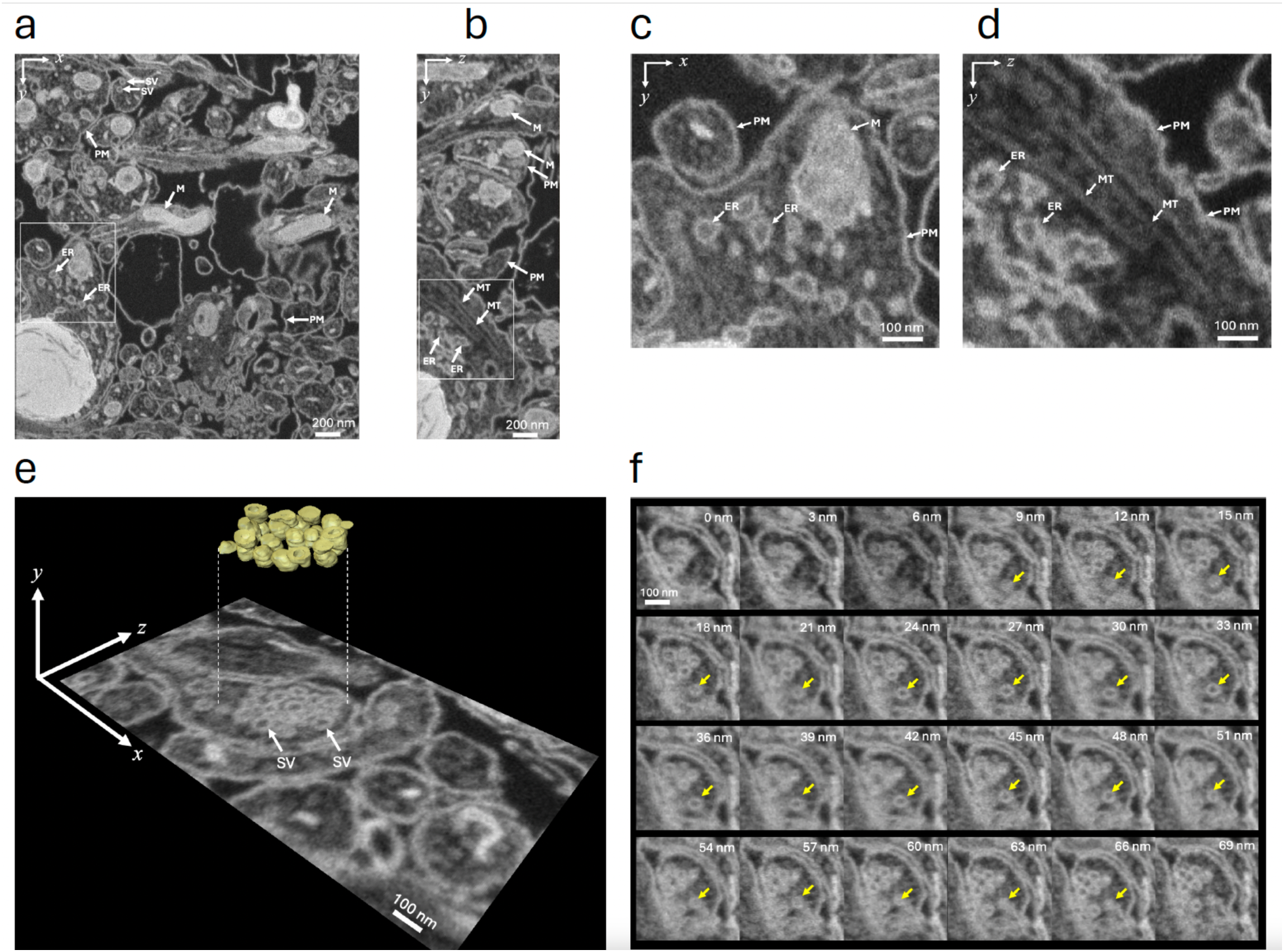
Orthoslices from 3D dataset acquired with the FIB-SEM from sample of three- week-old, cultured, dissociated, rat hippocampal neurons that were fixed and stained before embedding in Araldite/Epon, **(a-f).** Representative orthoslices in the *x*-y plane (**a) and in the *y*-*z* plane (**b**); PM, plasma membrane; SV, synaptic vesicle; M, mitochondrion; ER, endoplasmic reticulum; MT, microtubule. Higher magnification views of the ultrastructure in *x*-*y* plane (c) and *y*-*z* plane (**d**) indicate near-isotropic resolution of 3D FIB-SEM images. Slice in *x*-*z* plane (**e**) containing a presynaptic neuronal terminal with numerous synaptic vesicles (SV) of diameter 40±10 nm revealing lumen of vesicle, and 3D visualization in inset. Pixel size was 3 nm (in *x*, *y*, and *z*); pixel acquisition time was 16 µs for e^-^ beam; FIB Ga^+^ current was 700 pA, with concurrent scanning of the Ga^+^ and e^-^ beams. Examining the widths of membranes in the *x*-*y* plane in (a) indicates a resolution of approximately 5-10 nm, and in the *z*-direction a resolution of ∼10 nm. Series of *x*,*y* orthoslices spaced by 3 nm in region of a synaptic terminal (**f**) with the increment along the *z*-axis indicated for each slice; a single synaptic vesicle of diameter ∼50 nm, marked by the yellow arrows, is observed in 18 successive slices.**

Although we expect that some Ga^+^ ions will be deposited in the layer close to the block-face, it is not clear how many atoms are implanted in relation to atoms contained in the Epon/Araldite polymer embedding the biological structures together with bound heavy atoms of stain. In the present study, we show that the excellent performance of the FIB-SEM for 3D imaging of cells and tissues is due to an important feature of the focused Ga^+^ beam acting on epoxy-resin embedded specimen blocks of stained biological specimens. We have examined the FIB-SEM block-face using: (1) backscattered electron images to view a cross section of the block in the SBF-SEM; (2) high-angle annular dark-field (HAADF) images in the scanning transmission electron microscopy (STEM), as well as electron energy loss spectroscopy (EELS) data to obtain compositional information from FIB-milled sections of Epon/Araldite; (3) STEM-HAADF and STEM-EELS images from sections cut perpendicular to the FIB-SEM block-face to obtain compositional information from the region of Ga^+^ implantation; (4) we have performed Monte Carlo ion trajectory simulations to estimate the lateral spread of implanted Ga^+^ ions at the block face; as well as (5) Monte Carlo electron scattering simulations to model the backscattered electron signal from stained structures within the epoxy-embedded, gallium implanted tissue blocks.

## Results

### 1. Backscattered SBF-SEM images perpendicular to the block-face

We transferred specimen blocks that had been imaged in the FIB-SEM to a SBF-SEM, which enabled removal of sections perpendicular to the FIB-SEM block face using the built-in ultramicrotome. We first removed the protective Pt pad in addition to several microns of the Epon/Araldite block beneath the pad to expose a freshly cut cross section through the FIB-SEM block face, which was oriented perpendicular to the SBF-SEM block-face (as indicated in Fig. 1). At low magnification, several trenches are visible (Fig. 3a); backscattered electron images from these cross sections at higher magnification reveal a narrow line of width ∼25-30 nm at the edge of the block (Fig. 3b), indicating a composition with higher atomic number than the surrounding Epon/Araldite embedding resin. The material at the block face boundary sometimes appeared to be fractured suggesting a hardened layer. A line profile from the region indicated by the yellow band in Fig. 3b and averaged over a width of 200 nm shows a sharp peak in the BSE signal at the FIB-SEM block face, which provides our first evidence that there is a well-defined layer of Ga^+^ ion implantation. To investigate the gallium implantation further, we performed quantitative analyses using a 300-keV TEM-STEM, firstly on microtomed sections of the embedding material that had been milled with the Ga^+^ beam in the FIB-SEM, and secondly on sections directly cut through the FIB-milled block-face of pure Epon/Araldite blocks.

**Figure 3.**
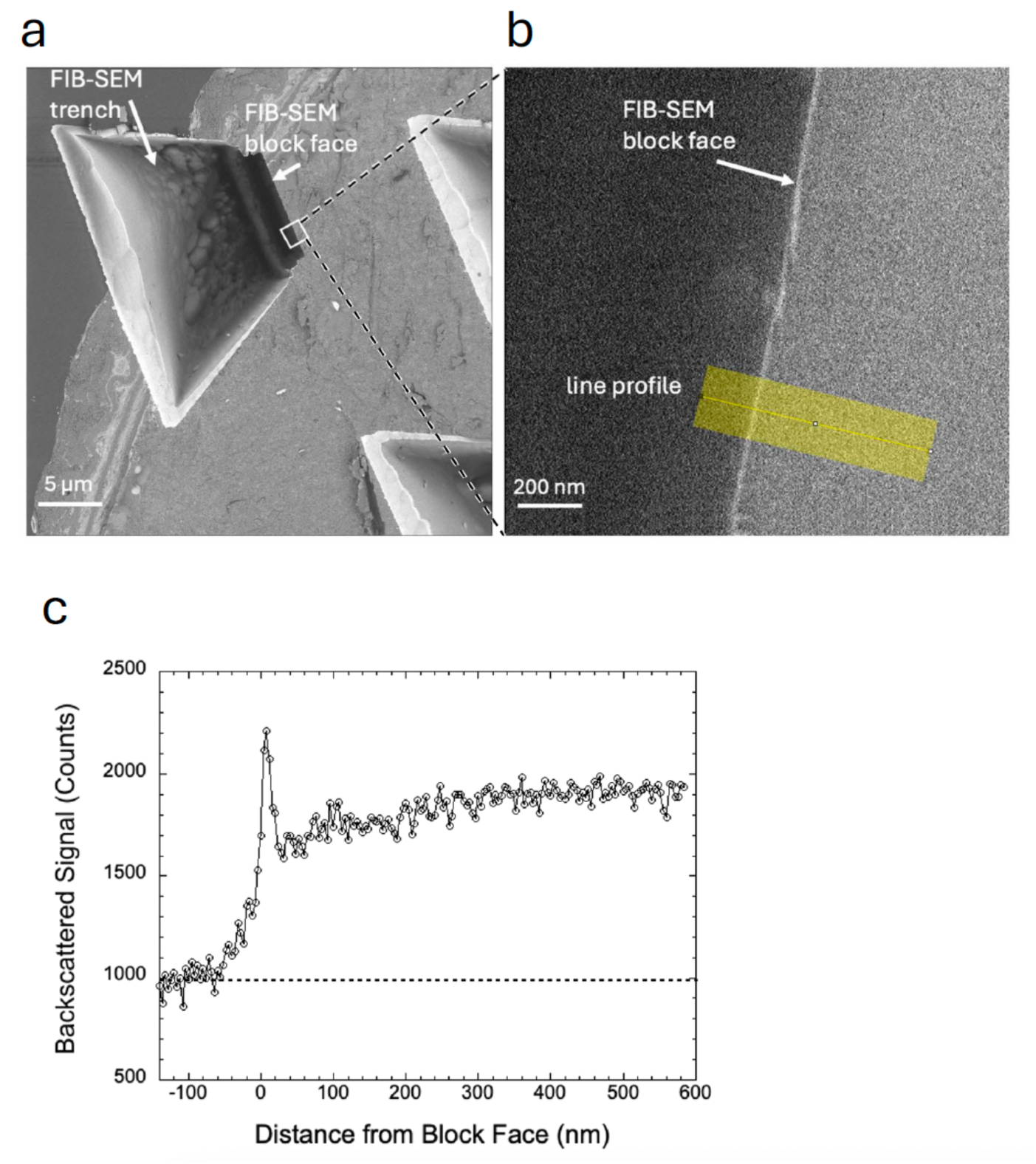
BSE images and line profile from FIB-SEM specimen block that had been transferred into an SBF-SEM. **(a,b,c)** Top view of trench at low magnification after removal of Pt pad by the in-situ diamond knife, showing location of straight block face at the edge of trench; the other three sides of the trapezoidal trench exhibit bright borders in the backscattered electron image even at low magnification indicating that gallium atoms are deposited at the high Ga^+^ current of up to 30 nA that is used to prepare the trench. This block also contains some heavy metal-stained biological tissue exhibiting contrast at the surface of the SBF-SEM block-face, which is perpendicular to the FIB-SEM block-face. A higher magnification view of the block face contained in the white square in (**a**) showing a bright line approximately 25 nm in width at the edge, and position of line profile (**b**). Some regions of the block face appear to be fractured. The line profile of the BSE signal averaged along a 200-nm width perpendicular to the block face shows a sharp increase in signal at the edge with a signal-to-background ratio of about 1:1 (**c**). To avoid the possibility that some of the contrast could be due to presence of heavy metal stain, the subsequent analyses in Figs. 4, 5 and Supplementary Fig. 5, 6, and 7 were performed on pure Epon specimen blocks.

### 2. STEM-HAADF and STEM-EELS analysis of FIB-milled sections of Epon/Araldite

A STEM high-angle annular dark-field (HAADF) image of a FIB-milled Epon/Araldite section microtomed to a nominal thickness of 200 nm is displayed in Fig. 4a, and the boxed subregion is shown at higher magnification in Fig. 4b, where a uniform intensity band of width ∼80 nm is evident at the milled edge. A line profile across the intensity band reveals a jump ratio >2 relative to the intensity of the Epon/Araldite away from the Ga^+^ FIB beam (Fig. 4c). If it is assumed that the C and O content of the Epon/Araldite remains unchanged in the FIB milling and the excess intensity above the polymer is due to Ga atoms, then based on the elastic scattering cross sections in Supplementary Table 3, the ratio of Ga:C is ∼0.10.

**Figure 4.**
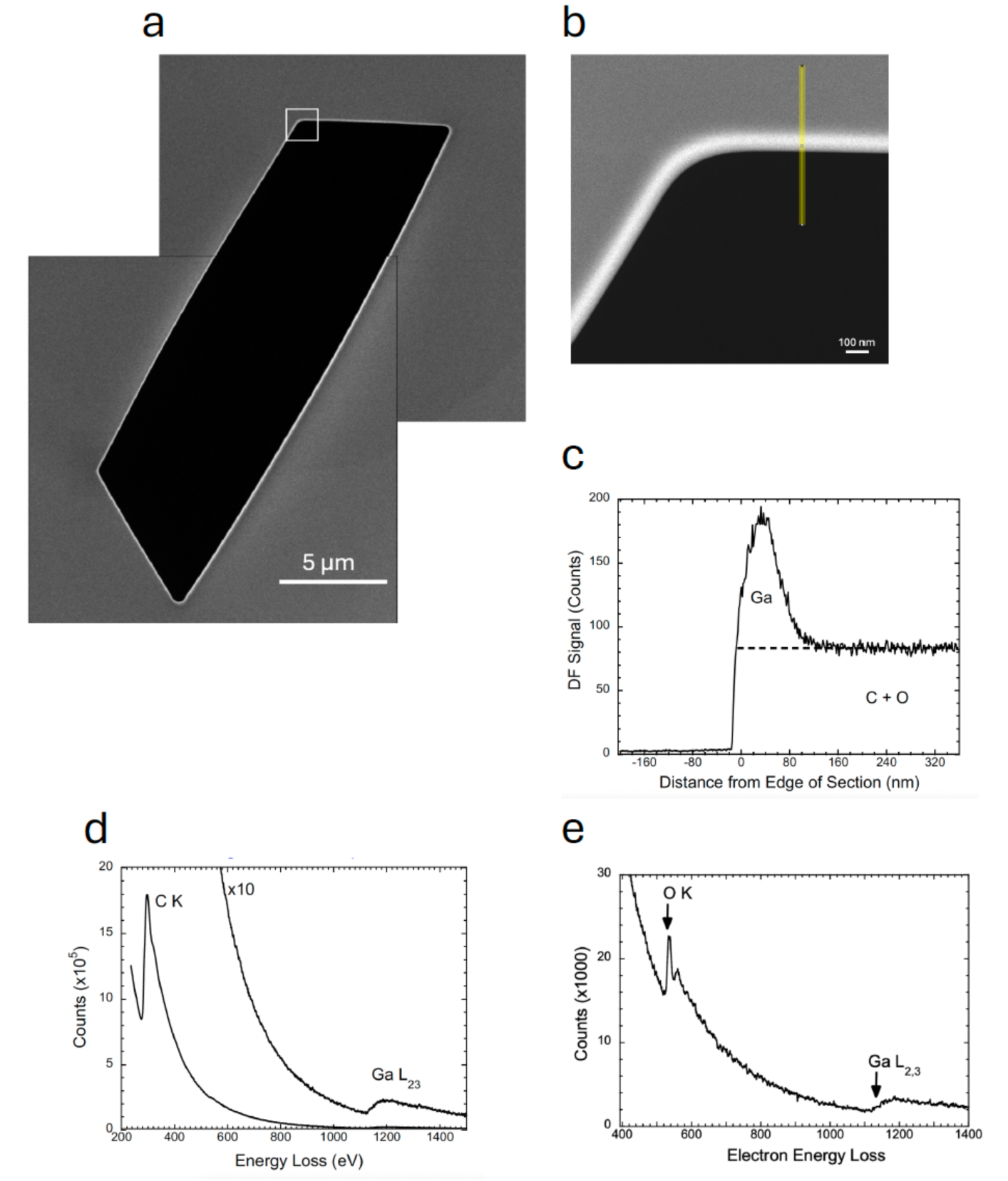
STEM-HAADF image of FIB-milled Epon/Araldite section cut to thickness of ∼200 nm with an ultramicrotome, together with line profile across the Ga^+^-milled edge and EEL spectrum from the region of higher HAADF intensity. **(a-e).** The HAADF image shows a higher intensity at the milled edge of the section in a low magnification image of the entire milled region of length ∼16 µm (**a**). A higher magnification HAADF image of the boxed region of interest in (a) reveals a uniformly increased intensity at the edge of the milled section (**b**), and a line profile shows a peak of width ∼80 nm with an intensity approximately twice that of the plastic section, which we attribute to increased elastic scattering due to implanted Ga^+^ ions; dashed line indicates extrapolation of HAADF signal from Epon/Araldite into region of implanted Ga^+^ ions (**c**). Energy loss spectrum shows presence of C K edge (285 eV) and Ga L2,3 edge (1115 eV) (**d**). Quantitative analysis was was performed using Ga2O3 particles deposited on a thin lacy carbon film as reference standard (**e**). The EELS analysis gave an estimated minimum Ga:C ratio of 0.117±0.012:1 using the method described in Supplementary Note 2.

In addition to the STEM-HAADF images, electron energy loss spectra (EELS) were acquired at a beam energy of 300 keV from the FIB-milled edges of the Epon/Araldite sections (Fig. 4d). Spectra integrated over areas of several hundred nm^2^ were quantified using reference spectra from Ga_2_O_3_ nanoparticles to determine the Ga L_2,3_ core shell cross section (Fig. 4e), and by measuring the EELS signal for the C K edge, Ga L_2,3_ edge, and O K edge (Supplementary Note 2). The energy window used for all inner shell edges was 100 eV and collection semi-angle was 10 mrad. The ratio of core edge cross section for the O K edge to the cross section for the C K edge was obtained from the EELS analysis plug-in of the Gatan Digital Micrograph software (Supplementary Fig. 1). Averaging EELS data from 10 regions of the milled edge of the Epon/Araldite section, we find an atomic Ga:C ratio of 0.117±0.012:1, which is consistent with the STEM-HAADF estimate of ∼0.10. In view of the high fluence of > 10^7^ e/nm^2^ required to perform the STEM-EELS measurements it is possible that some of the Ga atoms are displaced resulting in a lower Ga concentration than is evident in the HAADF images acquired at 4 orders of magnitude lower fluence of ∼10^3^ e^-^/nm^2^.

### 3. STEM-HAADF and STEM-EELS analysis of thin sections cut perpendicular to FIB-SEM block face

Next, we used an ultramicrotome to cut sections, of nominal thickness ∼200 nm, perpendicular to the Ga^+^ FIB-milled block faces in the regions of the Ga^+^ ion milled trenches. A schematic diagram of the sectioned block in Fig. 5a indicates a sectioned block of thickness 200 nm consisting mainly of carbon (green) with implanted gallium (red). STEM-HAADF images were acquired with a pixel size of 2 nm to 4 nm, as illustrated in Fig. 5b. A sharp increase in the STEM-HAADF signal is immediately evident at the edge of the sectioned FIB-SEM block face in Fig. 5b and the line profile across the block-face (Fig. 5c) shows a factor of >3 increase in the HAADF signal. The peak in the HAADF signal is very abrupt at the block face with a width of ∼6 nm, even though the nominal thickness of the section is more than a factor of 30 greater than this width. The peak, which is attributed to gallium implantation, has an asymmetrical shape with a tail extending ∼60 nm beyond the block-face, at which point the HAADF signal drops off to the level observed throughout the interior of the specimen block, i.e., far away from the FIB beam. This decay of the peak is suggestive of diffusion of Ga atoms into the block. The peak signal in the Ga layer has a mean jump ratio relative to the signal in the interior of the block of 3.3 ± 0.9, ranging from 2.3 to 4.2 (Supplementary Fig. 2). To check that the bright intensity at the sectioned block face is due to Ga^+^ implantation, rather than an artifact such as curling of the sections, a 200-nm section through a block that had not been FIB-milled (Fig. 5d) was imaged by STEM-HAADF (Fig. 5e); the line profile across the edge (Fig. 5f) shows a flat uniform intensity.

**Figure 5.**
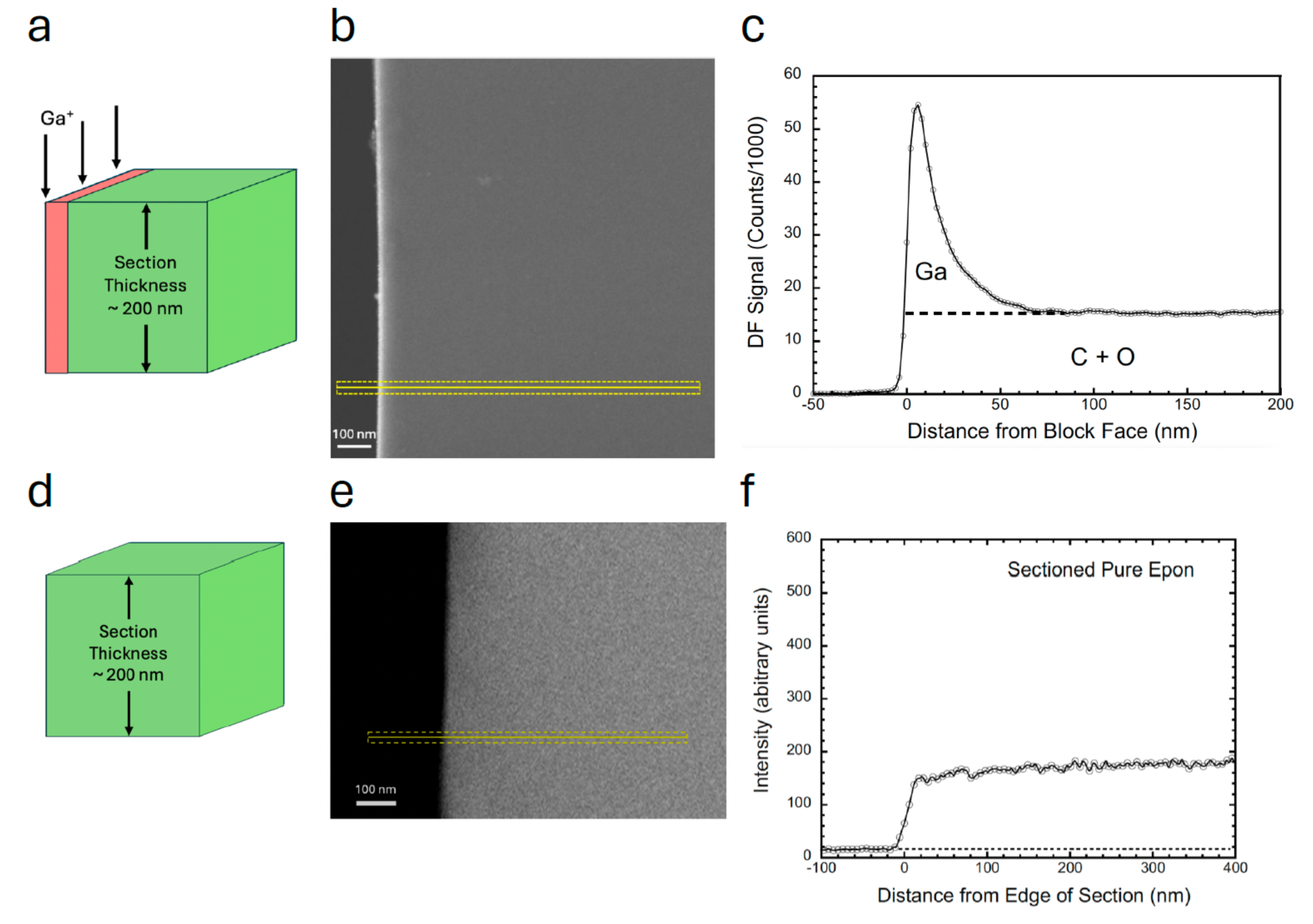
STEM-HAADF images of sectioned FIB-milled Epon/Araldite block face and unmilled block together with line profiles across the block faces, with both sections microtomed to a thickness of ∼200 nm. **(a-e).** Diagram of edge of sectioned block indicating narrow band of implanted Ga^+^ ions (**a**). HAADF image, acquired at beam energy of 300 keV with low electron fluence of ∼10^3^ e^-^ nm^-2^, shows a narrow bright band at the block face, which is attributable to Ga^+^ implantation (**b**). Intensity plot obtained across the block face in the region indicated and averaged over a 40-nm width parallel to the block face shows asymmetrical, sharp peak of width ∼25 nm with onset directly at the vacuum-block interface and a tail extending to ∼50 nm (**c**); plateau intensity far away from the Epon/Araldite block face corresponds to light atoms in the embedding material, mainly C and O, since the low elastic scattering cross section for H contributes negligibly to the HAADF contrast. Although it is difficult to ascribe the precise fraction of the DF signal that originates from C atoms at the edge of the FIB-milled block face because some atoms in the embedding material will have been sputtered away by the Ga^+^-FIB beam, we can estimate an upper limit to the number of C atoms as equal to number of C atoms in the undamaged embedding material (Supplementary Fig. 1, and Supplementary Note 1), as indicated by the dashed line in (c). To verify that there were no artifacts due to curling of the section at the block face, we imaged a 200-nm thick section of an Epon/Araldite block that had not been analyzed in the FIB-SEM, as illustrated (**d**). Neither the HAADF image in (**e**) nor the line profile along the normal to the block face in (**f**) show a peak in intensity at edge of the block.

In estimating the composition of the sectioned surface layer at the block-face from the elastic scattering signal, it is not necessary to consider H atoms because their contribution to elastic scattering is small, or to consider N atoms, which are present at low concentration (Supplementary Table 2). We do not know how many C and O atoms are lost from the region containing the strong Ga peak at the FIB-SEM block face, but it is clear from Fig. 5b that most of the observed signal at the surface layer of the milled block is due to scattering from Ga atoms. A lower limit for the ratio of gallium to carbon atoms in the surface layer can be obtained by extrapolating the dark-field line profiles from deeper in the FIB-SEM block, i.e., from the constant intensity region on the right side of the line profiles in Figure 5b and Supplementary Fig. 2, which has a value of: (*I_C_*)*_SL_* ≈ (*I_C_*_+0_)*_Epon_*. We know, however, that the undamaged epoxy resin (Epon) deeper inside the block contains approximately one-quarter the number of oxygen atoms as carbon atoms (Supplementary Tables 1 and 2), and we also know this oxygen is likely to be lost by a radiolytic interaction with electrons and Ga^+^ ions in the FIB-SEM. As discussed in Supplementary Note 1, the atomic ratio of Ga:C could be a factor of 1.36 higher than predicted if loss of oxygen is ignored, based on the elastic scattering cross sections of Salvat et al.^21^ and Jablonski et al.^22^ (Supplementary Table 3) and the composition of the Epon/Araldite embedding medium (Supplementary Table 2). If there is no loss of carbon from the region of high Ga implantation, the atomic ratio of Ga:C is 0.30±0.08:1, based on analyzing 6 STEM-HAADF images across the block-face in Supplementary Fig. 2. However, if oxygen atoms are lost by preferential sputtering of the Ga^+^ ion beam (or through radiolysis by the FIB-SEM’s electron beam), the Ga:C ratio could be as high as 0.41±0.11:1 and, if some carbon is also lost by sputtering from the surface layer of the block face that contains the high Ga concentration, the relative concentration of Ga at the block face could be even higher.

In addition to analysis of HAADF images, STEM-EELS images were also acquired from regions containing FIB-milled block faces, 7 of which are presented here, with one of these images being shown in Fig. 6a, and the six others in Supplementary Fig. 3. The top panel in Fig. 6a is the HAADF-STEM image, below which are the superposed C K-edge map (green) and Ga L_2,3_-edge map (red), showing co-localization of Ga atoms in the region with the strong HAADF signal, whereas the C map correlates with Epon/Araldite section, as expected; the two bottom panels are the separate Ga and C maps shown for clarity. To obtain sufficient Ga signal, the probe current was increased to ∼2 nA, the pixel size set to either 4.7 nm or 17 nm and the pixel dwell time from 0.1 s to 0.2 s resulting in a fluence of ∼5 × 10^7^ e^-^/nm^2^. Although the EELS data confirm the presence of substantial Ga^+^ ion implantation within ∼25 nm of the block-face, quantitative analysis using the method described in Supplementary Note 2 gave a ratio of Ga:C atoms that is substantially lower than the result obtained from HAADF-STEM images, which were acquired at a factor of >10^4^ lower electron fluence. The estimated EELS ratio of Ga:C was 0.038±0.007:1, a factor of ∼7 to 10 lower than the HAADF-STEM estimate. We believe that the discrepancy can be explained by radiation damage due to the high fluence of 300 keV electrons in the EELS measurements, including destabilization of the Ga-C glassy metastable phase by knock-on damage of carbon atoms in the implanted layer which could result in a carbon concentration that falls below a critical level. It is also possible that the high electron fluence contributes to the decrease in Ga:C ratio, by causing the ∼25 nm thin glassy layer to be tilted away from its vertical orientation, as illustrated in Supplementary Fig. 4.

**Figure 6.**
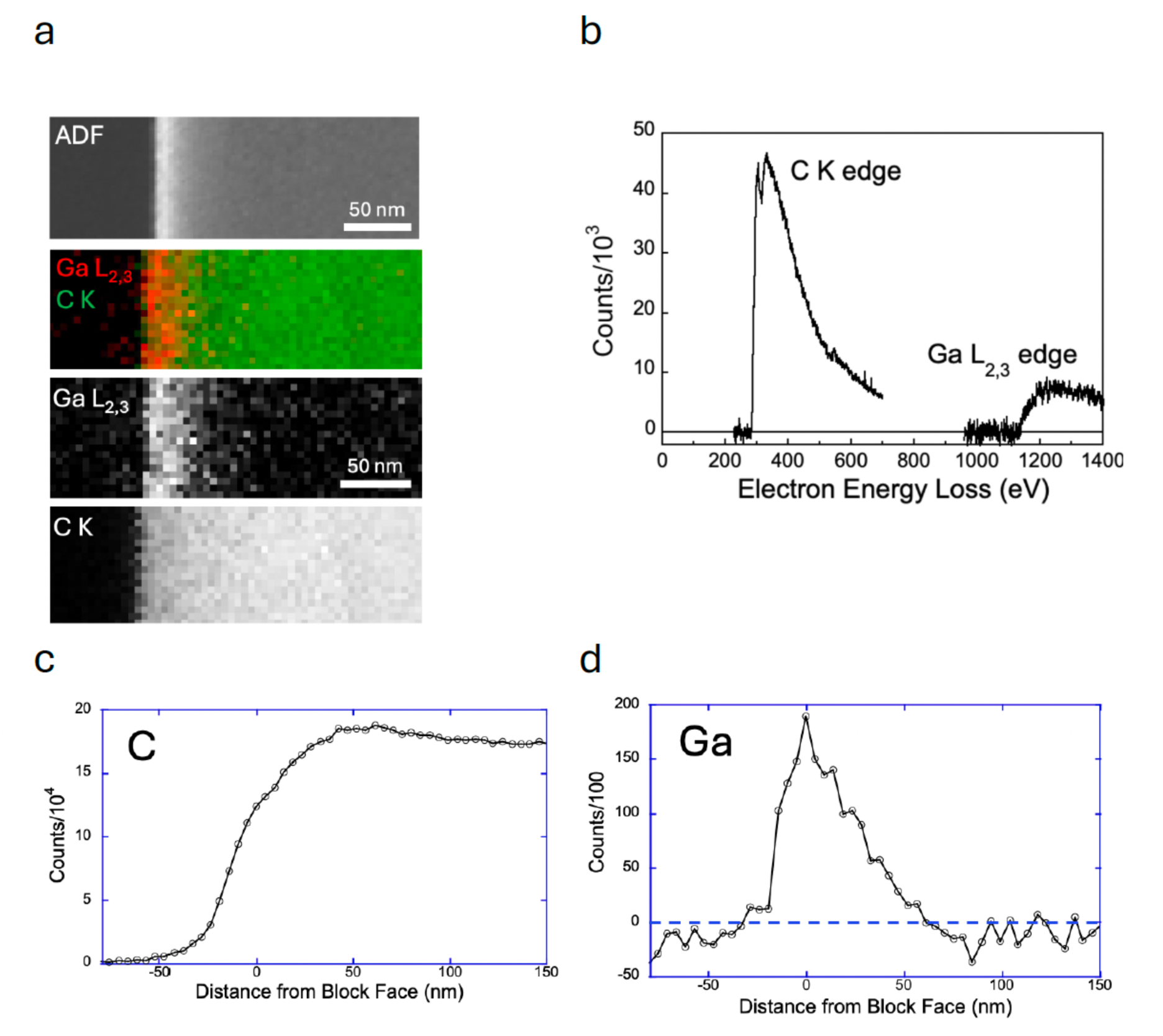
STEM-EELS compositional images from 200-nm thick section of FIB-milled Epon/Araldite block and EEL spectrum from the Ga-containing ∼25-nm outer layer of the milled block, together with C and Ga line profiles across the block face. **(a-d).** STEM-EELS data acquired from the region of the block face (**a**), where a layer of Ga (identified by the Ga L2,3- edge) corresponds to the region in the HAADF-STEM image showing elevated contrast, and a slight decrease in the C K-edge signal, possibly due to sputtering of C atoms by the Ga^+^ beam in the FIB-SEM or to knock-on damage by the 300 keV electron beam in the STEM-EELS measurement, where the electron fluence was >10^7^ e^-^ nm^-2^, i.e., 4 orders of magnitude higher than used for the HAADF-STEM measurements. EEL spectrum from the Ga^+^-implanted layer showing C K-edge and Ga L2,3-edge (**b**), as well as C and Ga EELS line profiles across the block face (**c**) and (**d**), respectively; line profiles show a broadening of the Ga peak at the block face from ∼25 nm to ∼50 nm, indicating that there could be beam-induced diffusion or escape of Ga atoms under the high electron fluence needed to collect the EELS compositional data.

Sometimes, we found that the sectioning of the FIB-milled block face became irregular, and this provided some interesting views of the hardened surface layer, as shown in Supplementary Fig. 5. Here, a HAADF STEM image acquired at a fluence of < 10^3^ e^-^ nm^-2^ shows that the Ga-rich glassy layer has fractured and rotated during cutting to produce a flake of overhanging hardened material that is considerably deeper than the 200-nm thickness of the Epon/Araldite section. In this particular region of the broken block face, rotation of the TEM tomography specimen holder by 35° brings the flake back to its vertical orientation, and a line-profile now exhibits a narrow and intense peak in the HAADF signal of width ∼30 nm, which is ten times stronger than the HAADF signal from the 200-nm thick Epon/Araldite section. This example illustrates the hardness and thinness of the surface layer.

### 4. Monte Carlo simulations of Ga^+^ ion trajectories in Epon/Araldite specimen block

We performed simulations of Ga^+^ ion beam scattering inside the specimen block based on the Stopping and Range of Ions in Matter (SRIM) model, developed by J.F. Ziegler et al.^20^ The downloaded version used in this study was SRIM-2013. The elemental composition of Epon/Araldite block was based on the data in Supplementary Table 2. Altogether, 10^5^ trajectories were modeled with the 30-keV Ga^+^ beam incident on a semi-infinite specimen block at a glancing incidence of 89° to the normal. The spread of Ga^+^ ions was visualized as a thin slice through the 3D trajectories. The trajectories provide information about the penetration of the ion beam in the FIB-SEM specimen as well the lateral spread of the ions as they are scattered normal to the FIB- SEM block face (Fig. 7a and 7b). The simulation result (Fig. 7b) shows that lateral spread into the specimen block in the *z*-direction is ∼25 nm, which was consistent with the thickness of the observed Ga layer detected by STEM-HAADF and STEM-EELS. The simulation also shows that the longitudinal spread of Ga^+^ ions is approximately 70 nm in the direction of the beam, which explains why in the experimental ion milling process the beam is scanned repeatedly along the *x*- axis to ensure that the surface layer of the Epon block is ablated to successively deeper distances along the y-axis, eventually reaching the trench at the bottom of the block face.

**Figure 7.**
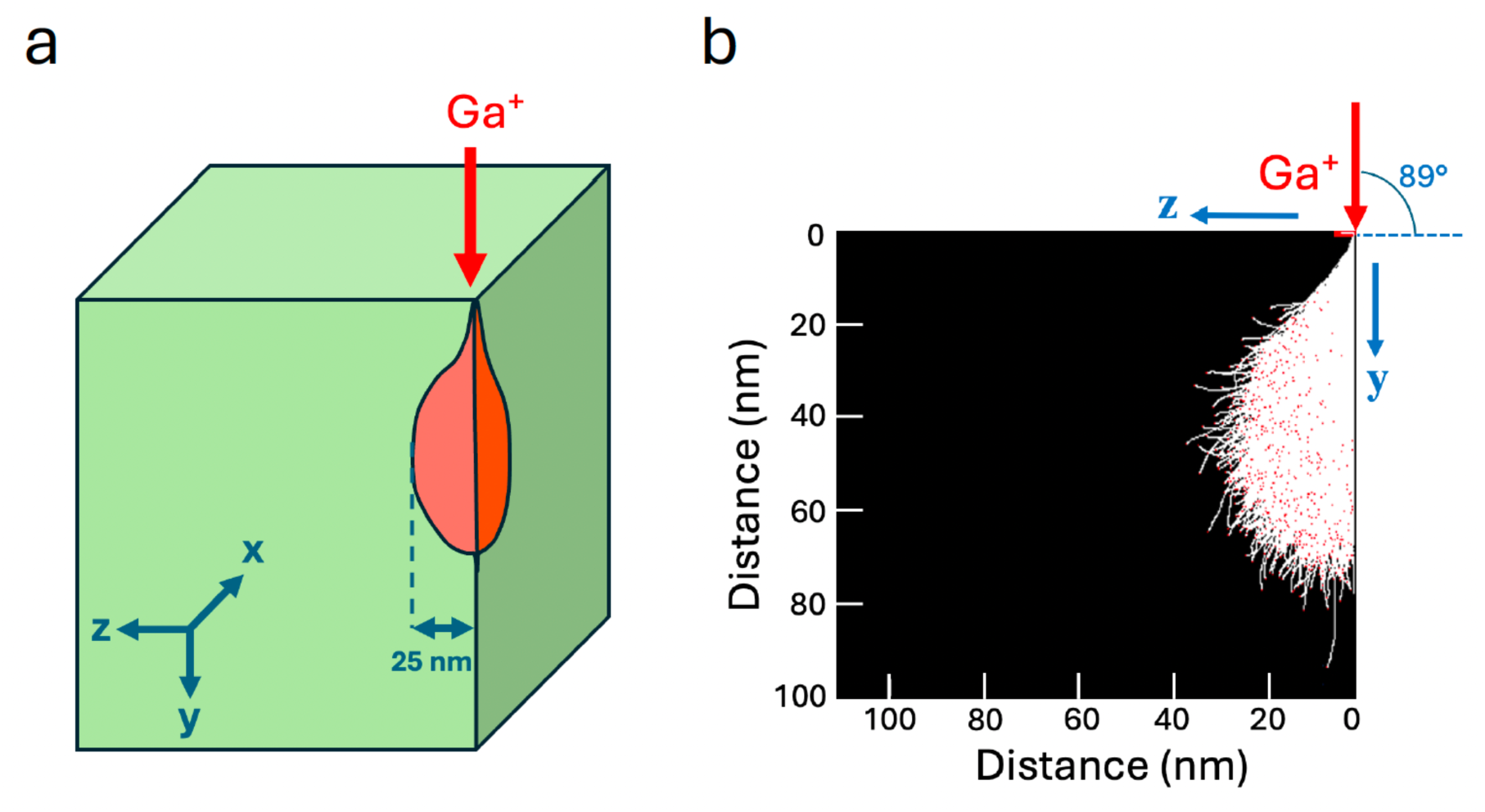
Simulated Ga^+^ ion trajectories at glancing incidence of 89° to the normal on a block-face of pure Epon/Araldite. **(a, b).** Schematic diagram showing incident Ga^+^ beam and coordinate system (**a**); Monte Carlo simulation of Ga^+^ ion beam scattering inside the block based on the Stopping and Range of Ions in Matter (SRIM) model and the Transport of Ions in Matter (TRIM) model (**b**). The composition of Epon/Araldite block was based on the elemental composition in Supplementary Table 2. Altogether, 10^5^ trajectories were modeled with a 30-keV Ga^+^ beam incident at a glancing angle of 1° to the *y*-axis of the specimen block (i.e., 89° to the normal to the block-face along the *z*-axis) as indicated. The glancing incidence angle of 1° allows scattering of Ga^+^ ions by the block, whereas the simulation breaks down with the glancing angle set to 0°. The lateral spread of Ga^+^ ions in the *z*-direction was visualized as a thin *y-z* slice in the *x*-direction through the 3D trajectories **(b)**. The lateral spread along the *z*-direction is ∼25 nm, which is consistent with the experimental data in Fig. 5 and 6.

**Figure 8.**
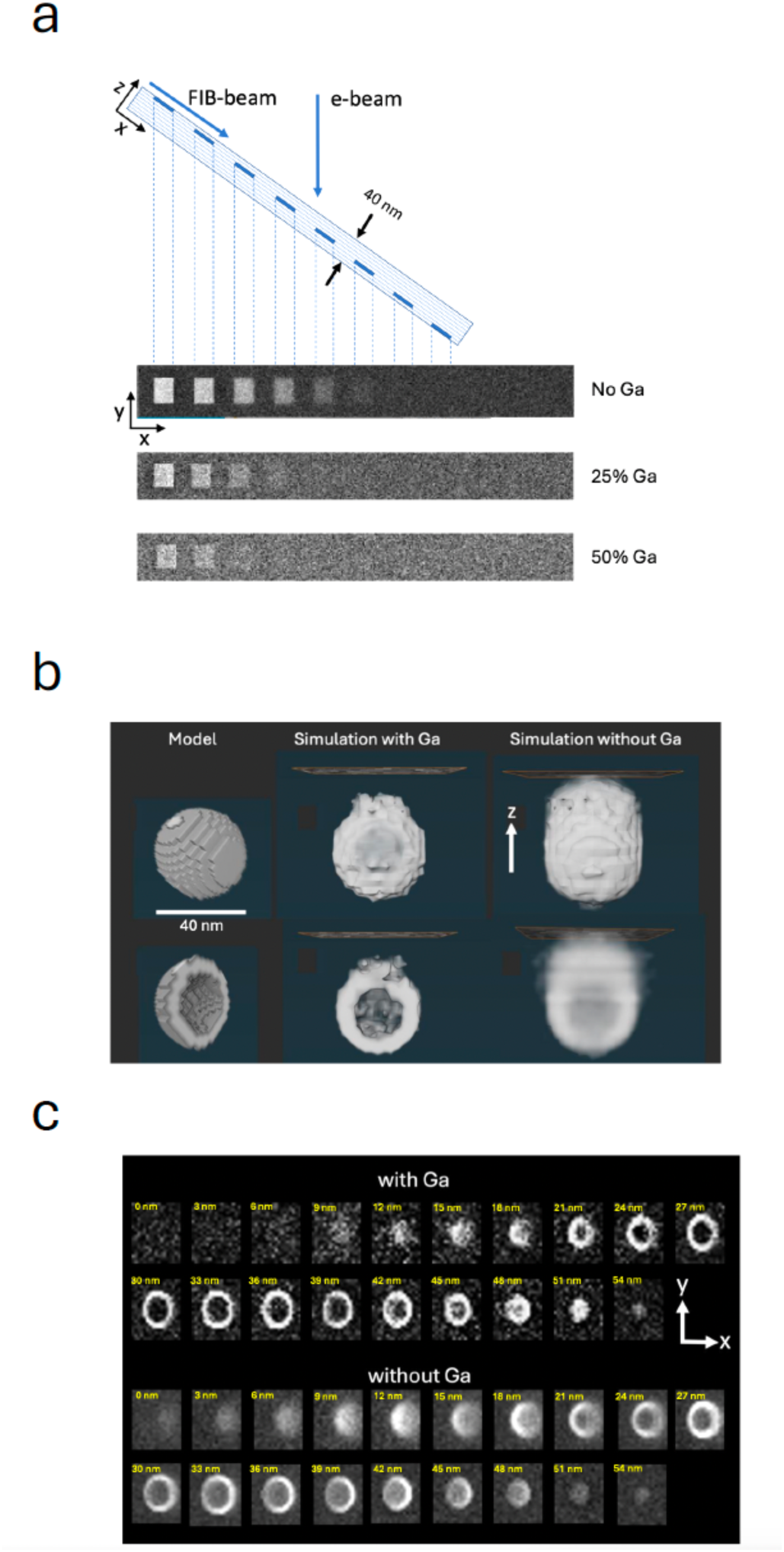
Simulated backscattered electron (BSE) images of heavy-atom-stained structures with and without Ga^+^ ion implantation (a, b, c). Monte Carlo simulations of backscattered electron images from regions of 3% atomic concentration of heavy metal (Pb) stain in an Epon/Araldite block using the software MC XRAY1.7. (a).^22^ Simulations were performed for 1.5 keV electron beam, and 3,000 trajectories per pixel. Contrast from regions of stain were computed for a block containing 25 atomic % Ga^+^, 50 atomic % Ga^+^, and 0% Ga^+^ implantation. Stain was located at depths of 0-5 nm, 5-10 nm, 10-15 nm, 15-20 nm, and 20-25 nm below the surface (a). With no Ga implantation, five regions are visible indicating a *z*-resolution of 20-25 nm; with 25% Ga implantation, three regions are visible indicating a *z*-resolution of 10-15 nm; and with 50% Ga implantation, two regions are visible, indicating a *z*-resolution of 5-10 nm. Monte Carlo simulation of the contrast of a single synaptic vesicle of diameter 40 nm with a membrane of thickness 5 nm (model) was computed for a Ga concentration of 50 atomic % and without Ga (**b**). Simulation with Ga predicts considerably improved resolution compared with simulation without Ga implantation. Simulated *x-y* orthoslices with and without Ga implantation are shown in **(c**). Without Ga implantation contrast is reduced substantially, and the stained vesicle is visible when situated below the surface, demonstrating loss of *z*-resolution. Voxel size was 3 nm × 3 nm × 3 nm, corresponding to the experimental conditions for images presented in Fig. 2.

### 5. Monte Carlo electron simulations of z-resolution in stained structures embedded in Epon/Araldite with and without Ga^+^ ion implantation

Having established from our HAADF-STEM measurements that the top 25 nm layer of FIB-SEM Epon/Araldite blocks contains high levels of gallium implantation, between ∼25 at. % and ∼50 at. % Ga, with the remaining atoms being mainly C, we ran Monte Carlo simulations to investigate the effect of the Ga on the quality of images obtained from stained biological structures using the software MC XRAY1.7. of Gauvin & Michaud^23^, based on earlier work by Hovington et al.^24^, Drouin et al.^25^, and Hennig & Denk^26^. We first considered a model structure consisting of Epon/Araldite in which 5-nm-thick square blocks of Pb at an atomic concentration of 3% were placed at different depths at thickness intervals of 5 nm. This concentration of approximately 3 atomic % of heavy elements associated with biological structures in specimens prepared for vEM^27^ had been previously estimated by He et al.^9^. The simulations were run (1) without any Ga implantation, (2) with 25 atomic % of Ga implantation, and (3) with 50 atomic % of Ga implantation. The stain was located at depths of 0-5 nm, 5-10 nm, 10-15 nm, 15-20 nm, 20-25 nm, 25-30 nm, 30-35 nm, and 35-40 nm below the surface. For each pixel in the simulated images 3,000 electron trajectories were computed. With no Ga implantation, five regions are visible indicating a *z*-resolution of 20-25 nm; with 25% Ga implantation, three regions are visible indicating a *z*-resolution of 10-15 nm; and with 50% Ga implantation, only two regions are visible, indicating a *z*-resolution of 5-10 nm (Fig. 8a).

In these simulations it was assumed that carbon atoms are replaced by gallium atoms. The results explain the previously puzzling finding that the *z*-resolution in FIB-SEM in the direction perpendicular to the block face is much higher than expected for primary electrons of energy 1.5 keV, which we use to acquire volume data in our laboratory. With 50 at. % gallium, the simulations show that the resolution is between 5 and 10 nm. Fig. 8a also reveals that although Ga reduces the contrast, there is sufficient signal-to-noise to detect the stained squares in the top 10 nm. Importantly, the simulation in the absence of Ga does not correspond to a real specimen because specimens imaged without Ga implantation (e.g., in SBF-SEM) collapse under electron fluences > 20 e^-^ nm^-2^. In fact, we have shown that samples analyzed by FIB-SEM can tolerate fluences > 10^4^ e^-^ nm^-2^. This resistance of FIB-SEM specimens to damage is consistent with the findings of Samira et al.^28^, who showed that Ga^+^ FIB beam interactions with epoxy resin could be used to microfabricate cantilevers for use in atomic force microscopy (AFM) and that these structures had a Young’s modulus of between 30 and 100 GPa, which is consistent with the observed exceptional stability of epoxy-embedded biological specimens that are imaged in the FIB-SEM.

We have found that FIB-SEM data sets allow us to visualize the 3D structure of individual synaptic vesicles in neuronal terminals, as illustrated in Figs. 2e, 2f, and in Supplementary Movie 1. Synaptic vesicles are spherical organelles of diameter ∼40 nm bounded by phospholipid bilayer membrane containing various membrane proteins, inside which are stored neurotransmitter molecules whose controlled release triggers neuronal activity. We have modeled the vesicle membrane as a shell of thickness 5 nm containing heavy atoms of stain at a concentration of 3 atomic %, as previously measured^9^, and we have performed Monte Carlo simulation of FIB-SEM image stacks with and without 50 atomic % Ga implantation, using a voxel size of 3 nm in *x*, *y*, and *z*. The model that includes Ga implantation predicts considerably greater spatial resolution than the simulation without Ga^+^ implantation (Fig. 8b). Simulated *x*-*y* slices for the modeled synaptic vesicle (Fig. 8c) at *z*-increments of 3 nm through the volume, again predict a higher spatial resolution with Ga^+^ implantation. Without Ga^+^ implantation the contrast is reduced substantially, and the stained vesicle becomes visible when it is situated below the surface. These results explain the improved *z*-resolution in FIB-SEM relative to the predicted z-resolution in the absence of a substantial level of Ga^+^ ion implantation.

## Discussion

Whereas users of FIB-SEMs have been able to obtain nanoscale 3D images of cellular ultrastructure that is almost isotropic, they have not fully understood the physical basis of why this capability is achievable and why such a high electron fluence can be delivered in the FIB-SEM without collapse of the specimen block. In SBF-SEM, the maximum electron fluence on the specimen is 100-1000 times lower than the fluence used in FIB-SEM. The higher electron fluence used routinely in FIB-SEM, which is required to obtain adequate signal-to-noise ratio with pixel sizes of 3-5 nm, would cause an intolerable collapse of the specimen if it had the same composition as the sample that is analyzed in the SBF-SEM, which would prevent acquisition of volumetric data. Yet the sample preparation is essentially the same for FIB-SEM and SBF-SEM. The only explanation for this apparent contradiction is that the sample in the FIB-SEM must undergo an extreme physical change, while preserving the delicate structures constituting the stained biological ultrastructure.

By analyzing the composition at the surface layer of the block face in stained biological samples embedded in epoxy resin, we can now understand how Ga^+^ FIB-SEM provides 3D ultrastructure of cells and tissues with a high, almost isotropic, spatial resolution as low as 5 nm. In the present study, we have discovered that the ion beam causes extremely high levels of gallium implantation reaching levels of up to 40 or 50 atomic % within a 25-nm wide layer of the surface. This finding, which has never been reported in the context of vEM of biological specimens, is consistent with the microfabrication of AFM cantilevers reported by Samira et al.^28^, and the formation of a glassy phase with a high Young’s modulus. (Hard skins on polymer samples formed by interaction with Ga^+^ beams were first observed by Moon et al.,^29^ without quantitative analysis of the implanted layer.) Our results indicate that a similar interaction occurs over the entire block face in FIB-SEM imaging of epoxy resin-embedded biological specimens, thereby converting the stained biological structures into a strong ∼25 nm layer of metastable Ga-C metallic glass that can withstand beam damage, yet retain the distribution of heavy-atom stain, thereby restricting penetration of the 1.5 keV electron probe into the subsurface layer. This transformation of the surface layer results in a *z*-resolution of 5-10 nm rather than the expected resolution of 20 to 25 nm if a much lower level of gallium were implanted. The higher tolerated electron fluence of >10^4^ e^-^/nm^2^ enables features in the images to be visualized, even though the intrinsic contrast is decreased by the gallium implantation; this is achievable because the signal-to-noise ratio is sufficient to detect features having reduced contrast. In addition, the gallium implantation increases the electrical conductivity of the entire block face preventing the specimen from charge build-up, even when individual cells are dispersed and surrounded by pure epoxy resin, which offers no electrical path to ground.

Since we have shown that it is possible to acquire backscattered electron images with a voxel size of 3 nm along the *z*-axis, we deduce that only a ∼3-nm thick outer zone of the ∼25-nm wide subsurface Ga-C metallic glassy phase is ablated under continuing exposure to the Ga^+^ FIB, in the time taken (∼60 s) to acquire a backscattered image from the entire block face. We suggest that continued exposure to the Ga^+^ ion beam in this outer surface layer sputters away C atoms dropping the carbon concentration to a critical value, below which the structure becomes unstable resulting in a phase transition with complete loss of material from the outer ∼3-nm surface layer, i.e., ablation. Meanwhile, additional scattered Ga atoms diffuse into the specimen block moving the ∼25 nm glassy phase further into the block, enabling collection of ultrastructural images from successively deeper layers underlying the initial block face.

We find that the previously accepted explanation for the high-spatial resolution perpendicular to the block-face in FIB-SEM is insufficient. It had been believed that energy-selection of the backscattered electrons from the specimen can restrict the range of depths from which the backscattered signal is generated by excluding multiple inelastic scattering. In many FIB-SEMs, including the one we have used in the present study, a voltage is applied to a grid placed in front of an energy-selected backscattered-electron (ESB) detector, which repels backscattered electrons with energies lower than the grid voltage. For example, typically the primary beam has an energy of 1,500 eV and a 700 V bias is applied to the grid. We acquired images of cultured neurons at different grid voltages, but we have found very little difference in resolution at the different grid voltages, even when no voltage is applied to the grid, as illustrated in our *x*-*z* plane orthoslices in Supplementary Fig. 6.

Our results further suggest that future biological FIB-SEMs using a plasma ion beam, e.g., He^+^, Ne^+^, Xe^+^ might not provide such a high *z*-resolution because of the lower average atomic number of the biological matrix in the absence of Ga^+^ ion implantation. Furthermore, the specimens are likely to be more susceptible to damage by the electron beam, as in the SBF-SEM, even though artefacts due to the gallium implantation are avoided^30^.

Finally, one might speculate that, had not the Ga^+^ FIB-SEM already been invented as a lift-out preparation technique for inorganic materials characterization by transmission electron microscopy, the same tool might have been designed specifically for vEM. The design criteria would include (1) reducing the range of distances below the surface of the epoxy resin-embedded specimen block from which low-energy (∼1.5 keV) backscattered electrons are generated by implanting a high concentration of an element with intermediate atomic number; (2) avoiding electric charge build-up by implanting a metallic element; and (3) converting the radiation- sensitive epoxy resin containing the heavy atoms of stain (i.e., Os, Pb, and U) into a strong, rigid, and radiation resistant inorganic glassy phase. One might expect that the optimal atomic number of the implanted ion should be roughly the geometric average of the atomic number of the light organic elements contained in the epoxy resin plus embedded biological material (*Z_org_*) and the atomic number of stain (*Z_stain_*). This would ensure that the electron range is reduced by a substantial factor, while also ensuring that the atoms of stain are still visible when the implanted ion has a high concentration. We might then expect that:

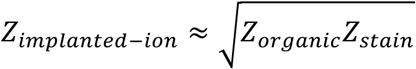

Taking a mean value of *Z_or_*_5*anic*_ = 7 (ignoring H atoms), and *Z_stain_* = 82 (the mean value for Os, Pb, and U atoms), we predict an optimal value for 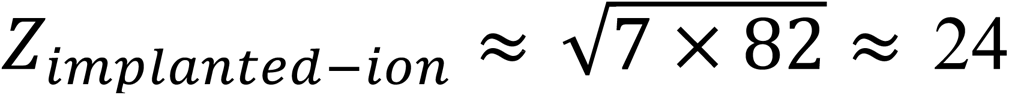, which is relatively close to *Z_Ga_* = 31. This choice of atomic number for the implanted ion thus ensures that the electron range is reduced by a substantial factor, while also ensuring that the atoms of stain are still visible above the background of the implanted ions. In this context, it is interesting to consider whether alternative liquid metal alloy ion sources (LMAISs) might potentially offer advantages over Ga^+^ ions sources.^31^ To achieve high spatial resolution in vEM, it is advantageous to use isotopically pure ion beams that can be focused to a single spot which is scanned parallel to the block face to mill away very thin layers. Multi-element LMAISs, such as Mg^+^/Co^+^/Pd^+^ alloy, Au^+^/Ge^+^/Bi^+^/ Li^+^ alloy, Au^+^/Ge^+^ alloy, Au^+^/Si^+^ alloy, or Ga^+^/Bi^+^/Li^+^ alloy, generally incorporate a crossed electric field/magnetic field mass filter (**E** × **B** Wien filter). These sources are therefore able to provide ions of a single mass that can be focused into a nanoscale spot. The findings of the present study suggest that the optimal choice of ion for FIB-SEM of stained embedded biological specimens should: (1) preserve backscattered electron image contrast between the heavy atoms of stain bound to the biological ultrastructure and ion-implanted plastic embedding material in the surface layer of the specimen block; and (2) form a glassy phase that is resistant to damage and collapse. It would be interesting to test Ge^+^-liquid metal sources to determine whether there are advantages relative to Ga^+^-sources.

The fact that the Ga^+^ FIB-SEM works so well for vEM of plastic embedded biological samples is fortuitous, since the original applications of Ga^+^ FIB-SEM involved imaging the ultrastructure of inorganic materials^32^, including preparation of lamellae for high-resolution TEM imaging based on the lift-out technique^33^. Applications were directed to metals, semiconductors, and ceramics, rather than to biological specimens. Unlike the case of inorganic materials, where Ga^+^ ion implantation is considered to be an artifact near the ablated surfaces, we have demonstrated that Ga^+^ ion implantation in vEM of biological specimens serves an essential role in modifying the specimen to improve the *z*-resolution, to eliminate charge build-up, and to greatly reduce radiation sensitivity. In fact, in its current form, FIB-SEM of embedded eukaryotic cells and tissues would not work without the Ga^+^ ion implantation. This result has not previously been appreciated.

## Methods

All research reported in this work was approved by and complies with the NINDS/NIDCD/NCCIH (NIH) Animal Care and Use Committee (ACUC).

### Sample Preparation

Dissociated rat hippocampal neurons were plated on glia cells pre-plated on acid cleaned and poly-lysine coated glass substrate in a custom growth medium at 35°C and 5% CO_2_ for 21 days before being fixed in 4% glutaraldehyde for 1 hr at RT. Samples were further fixed using 0.1 M cacodylate buffer containing 2.5% glutaraldehyde and 2 mM CaCl_2_ for 1 h in ice, and they were then washed three times with cold 0.1 M sodium cacodylate buffer containing 2 mM CaCl_2_ and spun at 600 × g for 5 min. Samples were then fixed in 3% potassium ferrocyanide in 0.3 M cacodylate buffer with 4 mM CaCl_2_ combined with an equal volume of 4% osmium tetroxide for 1 h in ice. After washing five times with H_2_O, samples were then placed in a 0.22 μm-Millipore-filtered 1% thiocarbohydrazide (TCH) solution in ddH_2_O for 20 min following five washes with double-distilled water (ddH_2_O) at RT each for 3 min. Samples were fixed in 2% osmium tetroxide in ddH_2_O for 30 min at RT following five washes with ddH_2_O at RT each for 3 min and then placed in 1% uranyl acetate (aqueous) for overnight at 4°C. The next day, samples were washed five times with ddH_2_O at RT each for 3 min and processed for *en bloc* Walton’s lead aspartate staining at 60°C for 30 min following five washes with ddH_2_O at RT each for 3 min. Samples were dehydrated before resin embedding. Blocks of Epon Embed-812 /Araldite-502 resin (Electron Microscopy Sciences, Hatfield, PA) having the same composition as blocks containing embedded biological samples in our laboratory, were prepared by mixing Embed-812 (25 mL), Araldite 502 (15 mL), DDSA (55 mL), BDMA (2.5 mL). The elemental composition of the embedding resin is provided in Supplementary Table 2.

### FIB-SEM

Specimens were imaged in a Crossbeam 550 focused ion beam scanning electron microscope (FIB-SEM) (Carl Zeiss Microscopy LLC, Thornwood, NY). A trapezoidal-shaped trench was ablated by the Ga^+^ beam (set to a high current of ∼30 nA) in front of the sample block with a narrower edge at the block face, i.e., at the location of the Pt pad, and a wider edge away from the block face (Figs. 1a and 1b) to allow access for the incoming 1.5 keV electron beam and the outgoing backscattered electrons from the block face. The block face was imaged with a 1.5 keV electron beam using an in-lens energy selective backscattered-electron (ESB detector) with the grid voltage set to 700 V to eliminate inelastically scattered electrons with multiple energy losses greater than 700 eV -- originating deeper in the specimen block -- from entering the detector. To illustrate the dependability of *z*-resolution on the grid voltage, we also acquired volumetric data at grid voltages of 0 V, 950 V, 1100 V, 1350 V, and 1500 V. Images were acquired with concurrent 700 pA electron and 700 pA Ga^+^ FIB beams, and with a pixel acquisition time of 16 µs. In addition to performing FIB-SEM imaging on Epon/Araldite blocks, which provided block faces of width ∼20 µm and height ∼20 µm, we also placed thin ∼200-nm ultramicrotomed slices of Epon/Araldite embedding material supported on EM grids into the stage of the FIB-SEM, tilted the grid to 54° to the horizontal and used the FIB to ablate the Epon/Araldite section using the same FIB current of 700 pA used to ablate the trench in the solid specimen block. Further details on the experimental set-up, important for understanding the rationale for the current study, are provided in the first section of the Results.

### SBF-SEM imaging

Specimen blocks that had been prepared in situ in the FIB-SEM (above) were removed from the FIB-SEM and transferred to a Zeiss Sigma VP SEM (Carl Zeiss Microscopy LLC, Thornwood, NY), which is equipped with an in-situ Gatan 3View (Gatan/Ametek, Warrendale, PA) serial block face ultramicrotome and an OnPoint backscattered electron detector, and a Carl Zeiss focal charge compensation (FCC) system to prevent specimen charging. The FIB-SEM block face was perpendicular to the surface of the specimen block (as indicated in Figs. 1a and 1b). The Pt pad was removed by cutting sections parallel to the surface of the specimen block. After the pad had been completely removed, the SBF-SEM block face contained the edge of the FIB-SEM block face. These images revealed an elevated backscattered electron signal at the edge.

### HAADF-STEM imaging

Specimen blocks that had been prepared in situ in the FIB-SEM (above) were removed from the FIB-SEM and transferred to a Leica EM UC6 ultramicrotome (Leica Microsystems, Buffalo Grove, IL). After removal of the Pt. pad from the top of the specimen block, sections of thickness ∼200 nm were cut through the trench and FIB-SEM block face. These sections were then deposited on square mesh copper EM grids. The sections were analyzed in an FEI/Thermo Fisher Tecnai TF30 TEM (Thermo Fisher Scientific, Hillsboro, OR) using dark-field STEM at a primary beam energy of 300 keV, and a Fischione Model 3000 high- angle annular dark field (HAADF) detector. The probe current was 160 pA with a pixel dwell time of 5 µs, and a nominal probe size of 2 nm, which gives an estimated electron fluence of 1.2 × 10^3^ e^-^ nm^-2^. Dark-field images were also acquired from 200-nm thick Epon/Araldite films that had been placed in the FIB-SEM and subjected to ablation by the Ga^+^ FIB beam.

### EELS analysis

EELS data were obtained from the 200-nm thick plastic sections that had been subjected to ablation by the FIB beam in the Zeiss Crossbeam 550 SEM. Electron energy loss spectra were acquired using the Gatan Digital Micrograph software in the STEM-EELS mode using the same Tecnai TF30 TEM used to collect the HAADF data. This instrument was also equipped with a Gatan Quantum Dual EELS Enfinium spectrometer/imaging filter (Gatan/Ametek Inc., Warrendale, PA). The beam energy was 300 keV. The energy loss range was 200 eV–1600 eV, and the elemental maps were taken from small areas of the specimen. To analyze the composition of the Ga^+^ FIB-milled edges of Epon/Araldite sections, EELS images containing about 400 pixels were acquired with a pixel size of 2 nm and a probe current of ∼1 nA, with an acquisition time of ∼0.1 s/pixel and an electron fluence of ∼10^8^ e^-^ nm^-2^. To analyze the composition of the sectioned Ga^+^ FIB-milled Epon/Araldite block, EELS images containing about 1000 pixels were acquired with a pixel size of 4.7 nm or 17 nm, and a probe current of ∼1 nA, with an acquisition time of 0.1 s to 0.2 s/pixel and an electron fluence of 5×10^7^ e^-^ nm^-2^. Electron energy loss spectra from the block face region were quantified using Digital Micrograph using reference spectra from gallium oxide, which were collected from Ga_2_O_3_ nanoparticles (Thermo-Fisher Chemicals, Waltham, MA) deposited on lacy carbon support films, and by analyzing the C K edge and the Ga L_2,3_ edge for the block face region, and the O K edge and the Ga L_2,3_ edge for the Ga_2_O_3_ nanoparticles. The energy window used for all inner shell edges was 100 eV and collection semi- angle was 10 mrad. The ratio of core edge cross section for the O K edge to the cross section for the C K edge (Supplementary Fig. 1) was obtained using Gatan Digital Micrograph version 3.21 with the EELS analysis plug-in^34^. EELS data were processed according to Supplementary Note 2.

### Simulations of Ga^+^ ion scattering in specimen block

Monte Carlo simulations of 30 keV Ga^+^ ions scattering were performed using the Stopping and Range of Ions in Matter (SRIM) software^20^ by downloading the SRIM-2013 program from the publicly available web site at http://www.srim.org/. Altogether, 10^5^ trajectories were modeled with the 30-keV Ga^+^ beam incident at a glancing angle of 89° to the normal on a semi-infinite specimen block of Epon/Araldite embedding material as the target with a known elemental composition in Supplementary Table 2. In the simulation, the lateral spread of Ga^+^ ions was visualized as the projection of a thin slice through the 3D trajectories. The simulation also provided the range of the Ga^+^ ions in the forward direction, i.e., parallel to the FIB-milled block face.

### Simulations of backscattered images

The MC X-Ray 1.7 Monte Carlo simulation software was used to simulate electron backscattered images from stained biological structures embedded in Epon/Araldite blocks with and without gallium implantation. The software, which was downloaded from https://montecarlomodeling.mcgill.ca/download/download.html, is in the public domain^22^. MC X-Ray is an extension of the Monte Carlo programs CASINO and Win X-Ray since it computes the complete x-ray spectra from the simulation of electron scattering in solids of various types of geometries. In this study we only used the software to generate backscattered electron images. We simulated the backscattered signal from square regions containing 3 atomic % Pb, which we have shown to have the highest concentration of the three heavy elements, Os (Z=76), Pb (Z=82), and U (Z=92), present in specimens prepared by UC San Diego NCMIR protocol that we use in our lab (He et al.^9^), and the atomic number of Pb is close to the mean atomic number of the three elements of stain. In the simulations, the regions of stain are 5 nm in thickness and are placed at different depths below the surface at intervals of 5 nm. The composition of the embedding material is given in Supplementary Tables 2 and 3. The simulations were carried out with no gallium implantation, with 25 atomic % Ga, and with 50 atomic % Ga following the experimental estimates of the possible ranges of Ga to C, based on the HAADF STEM and EELS data. Monte Carlo simulations were also performed on a modeled presynaptic vesicle of outer diameter 40 nm, with a membrane of thickness 5 nm and a stain concentration of 3%, with a voxel size in *x*, *y* and *z* of 3 nm × 3 nm × 3 nm. Simulations were performed for no Ga implantation and for 50% Ga implantation.

## Acknowledgments

The authors thank Ms. Christine Winters, Structural Cell Biology Section, National Institute of Neurological Disorders and Stroke (NINDS), NIH, for providing the specimen of cultured dissociated rat neurons.

## Author contributions

M.A.A. obtained the SBF-SEM images, STEM-HAADF images, and STEM-EELS images, performed the SRIM ion scattering Monte Carlo simulations, and analyzed the data; Z.Y. performed the backscattered electron Monte Carlo simulations, obtained FIB-SEM data and analyzed the data; J.C.K. obtained the FIB-SEM data, and analyzed the data; C.M.B. obtained FIB- SEM data, and analyzed the data; S.J.F. participated in the analysis and visualization of the data; X.C. prepared the specimens of cultured dissociated rat hippocampal neurons; G.Z. performed the specimen preparation for the FIB-SEM, SBF-SEM, and STEM; R.D.L. conceived the project, analyzed the data, and wrote the manuscript.

## Additional information

Supplementary information is appended to this paper.

