## Supplemental Information (Tables, Figures, and Notes) for "Biological volume-EM with a focused Ga^+^ beam depends on forming radiation- resistant Ga-rich layer at block face"

[*]Co-first author (equal contributions)

[**]Corresponding Author

#### Description of Additional Supplementary Files

##### Supplementary Tables
Table 1
Table 2
Table 3
Table 4

##### Supplementary Figures
Figure 1
Figure 2
Figure 3
Figure 4
Figure 5
Figure 6

##### Supplementary Notes

Note 1
Note 2

##### Supplementary Movies

Movie 1

#### Supplementary Tables

**Supplementary Table 1.** Composition of Epon/Araldite embedding resin used to prepare the samples. The density of the cured resin Epon/Araldite is 1.20±0.05 g/cm$^3$.

| Component | Mass used in mixture (g) | Formula unit | Molecular weight |
|---|---|---|---|
| EMbed 812 | 27.5 | $C_{12}O_6H_{20}$ | 260 |
| Araldite 502 | 17.0 | $C_{18}O_2H_{19}$ | 267 |
| Dodecenyl succinic anhydride (DDSA) | 55.0 | $C_{16}O_3H_{19}$ | 266 |
| Benzyl dimethylamine (BDMA) | 2.15 | $C_9NH_{13}$ | 135 |

**Supplementary Table 2.** Number density of elements in Epon/Araldite embedding resin, containing four chemical components (Embed 812, Araldite 502, DDSA, and BDMA), based on a resin density of 1.20 g/cm$^3$.

| **Element** | **Symbol in Supplementary Note 2** | **Number density (atoms/nm$^3$)** |
|---|---|---|
| C | $n_{\mathbf{C}}$ | 42.0 |
| N | $n_{\mathbf{N}}$ | 0.1 |
| O | $n_{\mathbf{O}}$ | 9.9 |
| H | $n_{\mathbf{H}}$ | 63.8 |

**Supplementary Table 3.** Elastic scattering cross section per atom ($\sigma_e$) for $E_0$ = 300 keV electrons collected by HAADF detector with inner angle $\alpha_{min}$ = $10^{-2}$ rad, and outer angle $\alpha_{max}$ = 5 x $10^{-2}$ rad. Data taken from Jablonski et al.[21]. The cross sections are given in units of the Bohr radius squared, $a_0^2$ = 2.800 x $10^{-21}$ $m^2$. These elastic scattering cross sections were used to characterize the composition of the sectioned FIB-milled block face in a 300 keV TEM/STEM (Thermo-FEI TF30) equipped with a Fischione Model 3000 HAADF detector to collect annular dark-field STEM images. See Supplementary Note 1.

| **Element** | **Symbol in Supplementary Note 1** | **Elastic scattering cross section per atom ($\sigma_e$) (units of $a_0^2$)** |
|---|---|---|
| C | $(\sigma_e)_C$ | 6.01 x $10^{-3}$ |
| N | $(\sigma_e)_N$ | 7.70 x $10^{-3}$ |
| O | $(\sigma_e)_O$ | 9.13 x $10^{-3}$ |
| Ga | $(\sigma_e)_{Ga}$ | 67.2 x $10^{-3}$ |

**Supplementary Table 4.** Inelastic scattering cross section per atom $\sigma_X$ for $E_0$ = 300 keV electrons with a collection semi-angle $\alpha$ = $10^{-2}$ rad, and energy integration window $\Delta$ = 100 eV for exciting atomic inner shell of element X. These inelastic inner-shell scattering cross sections were used to characterize the composition of the sectioned FIB-milled block face in a 300 keV TEM/STEM (Thermo-FEI TF30), equipped with a Gatan Quantum Model 965 Enfinium DualEELS electron energy loss spectrum-imaging system. See Supplementary Note 2.

| **Element** | **Inner Shell Excitation** | **Energy loss (eV)** | **Symbol in Supplemental Note 2** | **Inelastic scattering cross section ($\sigma_X$) ($cm^2$ per atom)** | **Computation or Measurement** |
|---|---|---|---|---|---|
| Carbon | K-edge | 285 eV | $\sigma_C(\alpha, \Delta)$ | $4.49 \times 10^{-21}$ | Gatan DMS v. 3.21 w/ EELS plug-in[34] |
| Oxygen | K-edge | 532 eV | $\sigma_O(\alpha, \Delta)$ | $1.26 \times 10^{-21}$ | Gatan DMS v. 3.21 w/ EELS plug-in[34] |
| Gallium | $L_{2,3}$-edge | 1,116 eV | $\sigma_{Ga}(\alpha, \Delta)$ | $0.53 \times 10^{-21}$ | EELS analysis of $Ga_2O_3$ nanoparticles |

### Supplementary Figures

#### Supplementary Figure 1

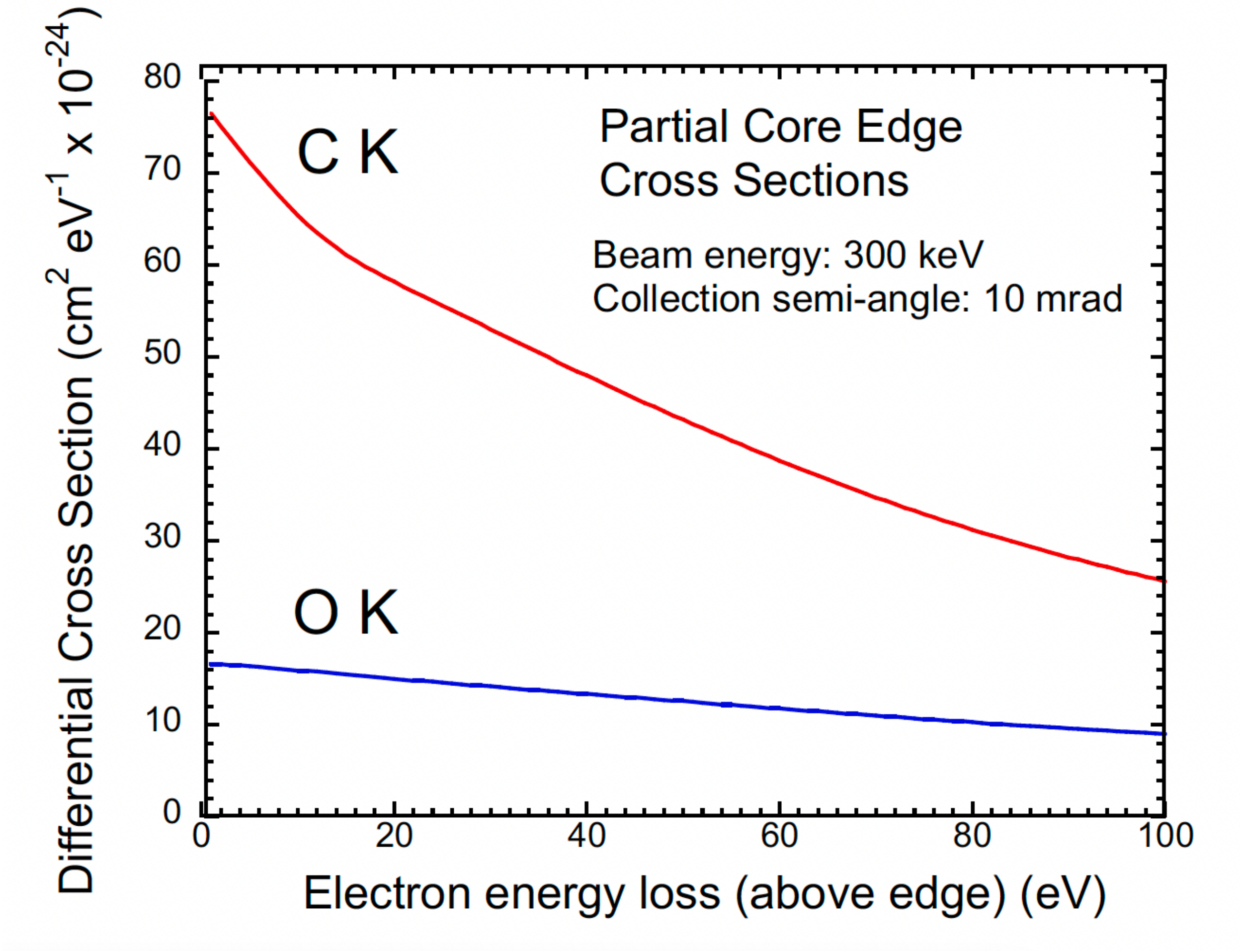


**Supplementary Figure 1 | Integrated EELS C K and O K edge cross sections per atom from the core edge threshold energies of 285 eV and 532 eV, respectively, up to Δ = 100 eV (integration window) above the edges, for an electron beam energy of 300 keV and collection semi-angle of α = 10 mrad.** Since the two edges are relatively close in energy, the ratio of cross sections for carbon to oxygen $\frac{\sigma_C\,(\Delta,\alpha)}{\sigma_O\,(\Delta,\alpha)}$ computed from Gatan Digital Micrograph EELS analysis module will be less sensitive to uncertainties in the collection semi-angle α.

**Supplementary Figure 2**

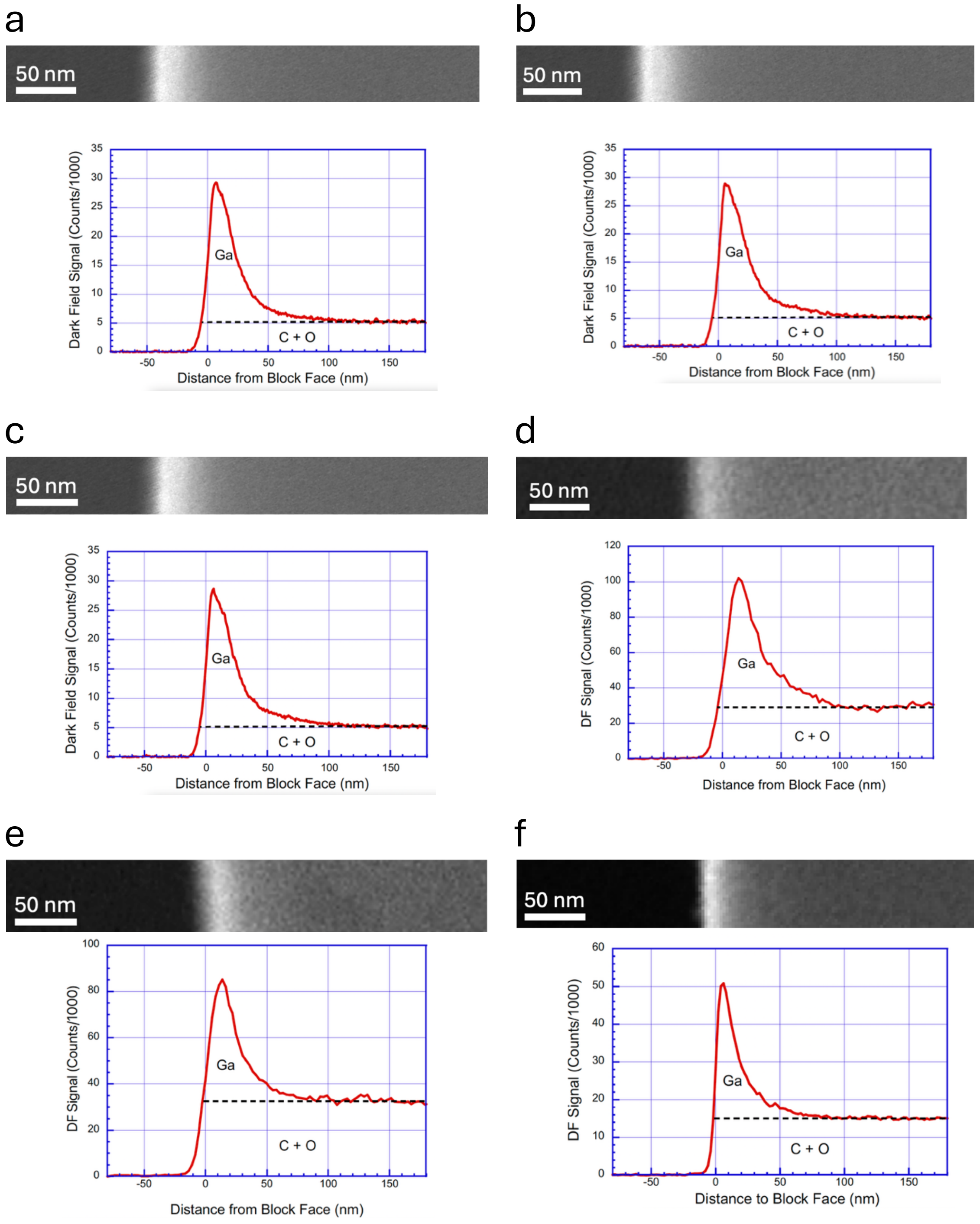


**Supplementary Figure 2 | Further examples of dark-field STEM images and line profiles across ~200-nm sections of FIB-milled Epon blocks revealing Ga-rich surface layer at the block faces (a,b,c,d,e,f).** Each image acquired at a beam energy of 300 keV shows a sharp 25 to 35 nm-wide peak in the HAADF intensity at the milled block edges, which is attributed to increased elastic scattering from the heavier implanted Ga atoms; further into the block the intensity decreases to a constant value corresponding to the elastic scattering from the epoxy resin (Epon) sections, containing mainly C and O as indicated in Supplementary Tables 1 and 2. By extrapolating the intensity of the Epon into the Ga-rich surface layer and subtracting from the peak intensity, a lower estimate of the Ga signal can be obtained. Then by using the computed elastic scattering cross sections for 300-keV electrons (Supplementary Table 3), we can estimate the Ga concentration.

#### Supplementary Figure 3

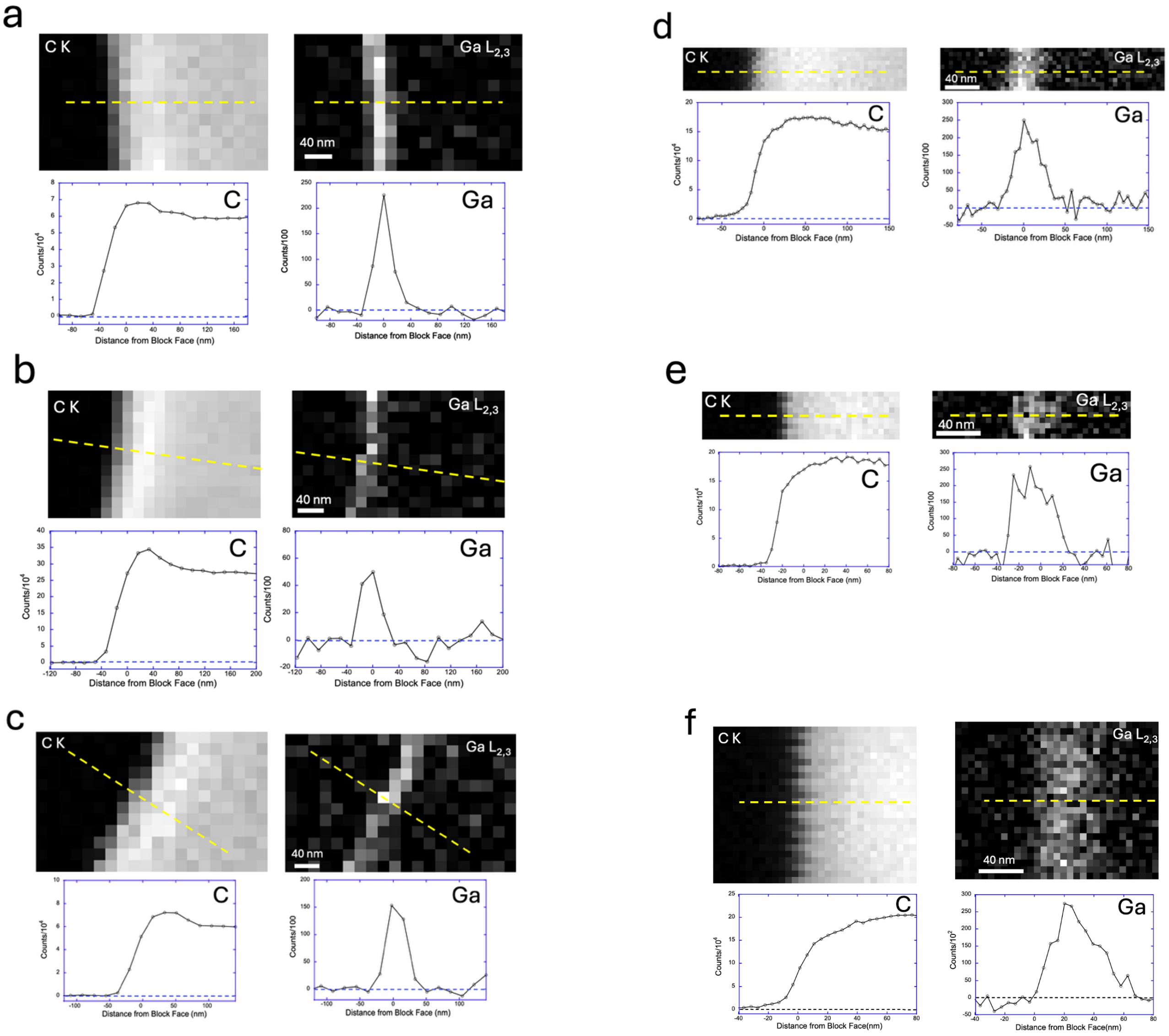


**Supplementary Figure 3 | Further examples of compositional maps of hardened Ga-rich surface layer.** Six electron energy loss (EEL) spectrum-images of 200-nm thick microtomed sections from $Ga^{+}$ FIB-milled Epon block in regions of the block face showing maps of C (K-edge) and Ga ($L_{2,3}$-edge), as well as line profiles along the dashed lines with pixel sizes of 17 nm and width of line profile along block face 200 nm **(a-c)**; and with pixel sizes of 4.7 nm and width of line profile along block face 100 nm **(d-f)**. Electron fluence was ~$10^{7}$ $e^{-}$ $nm^{-2}$ in (a-c), and ~$10^{8}$ $e^{-}$ $nm^{-2}$ in (d-f). Although these EELS maps show a localized layer of gallium at the sectioned FIB-milled block-face, quantitative analysis of 6 spectrum-images gave a Ga:C atomic ratio of only 0.038 ± 0.007, which is a factor of 7 to 10 lower than obtained by analyzing the dark-field STEM images acquired at much low electron fluence (e.g., in Fig. 5b). This discrepancy is attributed to the relative instability of the Ga-implanted block face after irradiation with the high fluence of 300-keV electrons that is required to perform EELS analysis at the Ga $L_{2,3}$-edge. Since Ga does not normally form carbides and gallium metal is a liquid at 30° C, loss of carbon by knock-on sputtering of the high fluence 300 keV electron beam could cause loss of cohesion of the metastable glassy phase, resulting in removal of gallium atoms. An apparent loss of Ga could also be caused by rotation of the partially detached surface layer in the sectioned block, as shown in Supplementary Fig. 4.

**Supplementary Figure 4**

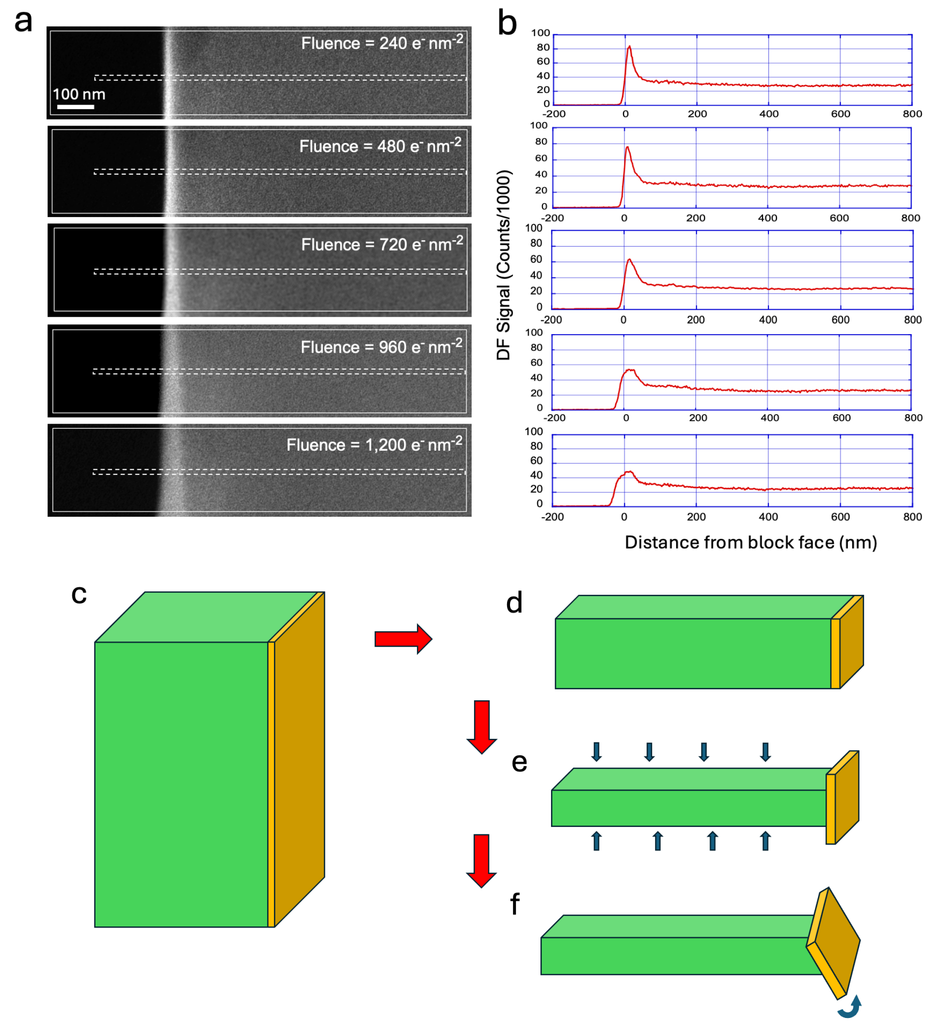


**Supplementary Figure 4 | Series of dark-field STEM images acquired at a beam energy of 300 keV from a 200-nm thick microtomed section through a $Ga^+$ FIB-milled Epon block in a region of the block face.** Each image was acquired at a fluence of 240 $e^-$ $nm^{-2}$, so that the fifth image had an accumulated fluence of 1,200 $e^-$ $nm^{-2}$ **(a)**. Although the first (top) image shows a narrow ~20 nm wide Ga-rich layer, damage of the specimen causes the sectioned layer to partially detach from the Epon and rotate, resulting in a width of ~80 nm and a corresponding weaker DF image intensity, which is also evident by the line-profiles across the images **(b)**. The tilting of the partially detached, sectioned, block-face layers is often observed in the dark-field images even at relatively low electron fluence ($<10^3$ $e^-$ $nm^{-2}$), whereas other DF images (e.g., in Fig. 5b and Supplementary Fig. 2) do not show detachment and rotation. This is one of the reasons why the experimentally measured Ga:C atomic ratio obtained from EELS is lower than expected based on the quantitative analysis of STEM dark-field images, since EELS analysis required several orders of magnitude higher electron fluence of $>10^7$ $e^-$ $nm^{-2}$. Schematic diagram of the Epon block (green) with a hardened Ga-rich glassy phase (yellow) **(c)**; sectioned block **(d)**; exposure of section to 300 keV electrons in STEM resulting in some thinning of the Epon even at a low electron fluence **(e)**; detachment and rotation of the surface layer **(f)**.

**Supplementary Figure 5**

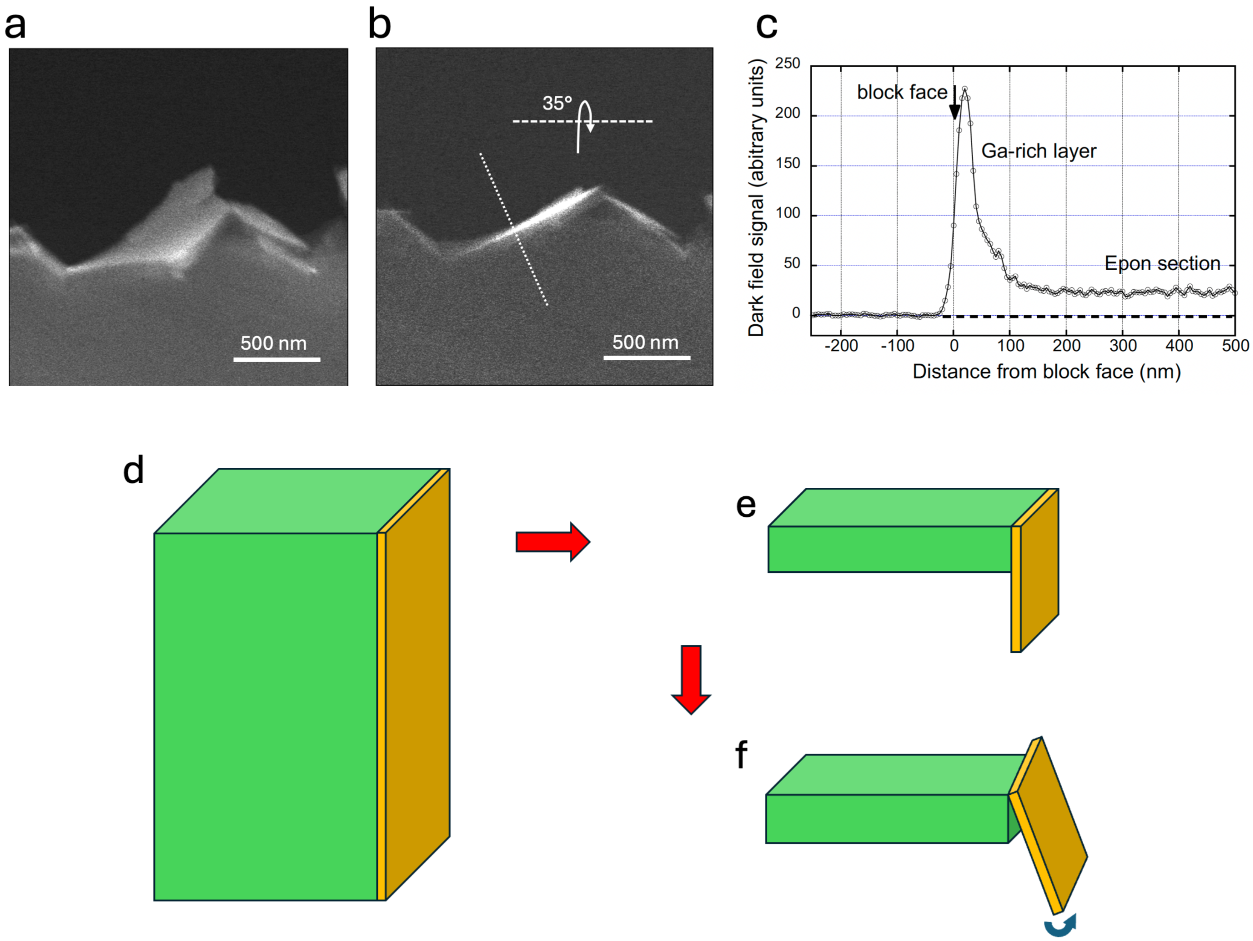


**Supplementary Figure 5 | Dark-field STEM images acquired at a beam energy of 300 keV from 200-nm thick microtomed section through a $Ga^+$ FIB-milled Epon block.** An image acquired at a fluence of < $10^3$ $e^-$ $nm^{-2}$ shows that the Ga-rich glassy layer has fractured and rotated during cutting to produce a flake of hardened material that is deeper than the 200-nm thickness of the Epon section, clearly demonstrating the presence of the distinct Ga-implanted surface layer **(a)**. Rotation of the specimen by 35° brings the flakes back to their vertical alignment **(b)**; and a line-profile across the image along the dotted line perpendicular to the vertically oriented flake shows a narrow and intense peak of width ~30 nm, which is ten times stronger than the intensity from the 200-nm thick Epon section **(c)**. Schematic diagram of the Epon block (green) with a hardened Ga-rich glassy phase (yellow) **(d)**; sectioned block with overhanging flake **(e)**; and rotation of the overhanging surface layer **(f)**.

**Supplementary Figure 6**

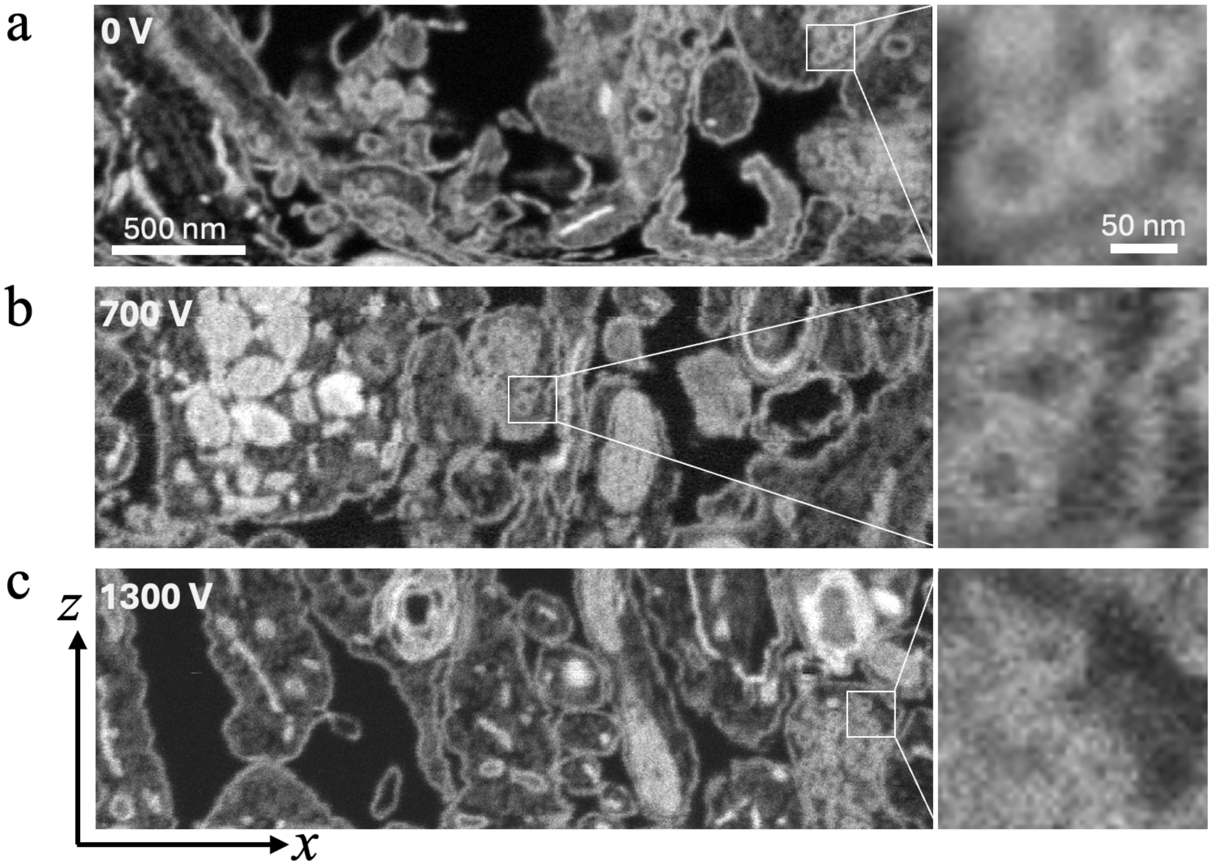


**Supplementary Figure 6 | Effect of energy selected backscattered grid voltage on z-resolution in FIB-SEM images.** Orthoslices in x-z plane for ESB detector grid voltage of 0 V (**a**); grid voltage of 700 V (**b**); and grid voltage of 1300 V (**c**). Clusters of synaptic vesicles indicated by boxes are visible regardless of grid voltage, with each vesicle containing a well-defined lumen surrounded by a membrane of width ~10 nm (expanded views of boxed regions). These images show that applying a grid voltage in front of the ESB detector to reject backscattered electrons, which have suffered inelastic scattering events deeper in the specimen block, does not account for high z-resolution in planes perpendicular to the block-face.

### Supplementary Notes

**Supplementary Note 1: Estimated composition of the FIB-milled surface layer of the Epon block, based on HAADF STEM images of sections cut perpendicular to the block face.**

The elemental composition of Epon/Araldite used to prepare the samples in this work is given in Supplementary Table 1. The cured resin has C, N, O and H atoms with composition $C_{42}O_{9.9}N_{0.1}H_{64}$, where the ratios represent the number of atoms per $nm^3$ (Supplementary Table 2). In estimating the elastic scattering signal, it is not necessary to consider H atoms because their contribution to elastic scattering is small or to consider N atoms that are present at low concentrations.

The HAADF-STEM image signal can be expressed in terms of the number densities of the atoms, and their elastic scattering cross sections at the 300 keV STEM beam energy, with the 10-mrad inner angle subtended by the HAADF detector, and the 50-mrad outer angle subtended by the HAADF detector. Denoting the elastic scattering cross sections for C, O and Ga by $(\sigma_e)_C$, $(\sigma_e)_O$ and $(\sigma_e)_{Ga}$, respectively, and the corresponding atomic number densities for C and O atoms in the Epon/Araldite by $(n_{C+O})_{Epon} = (n_C)_{Epon} + (n_O)_{Epon}$ and the number density of Ga atoms implanted into the surface layer by $(n_{Ga})_{SL}$, we can write:

$$\frac{(n_{Ga})_{SL}}{(n_C)_{Epon}} \approx \left[\frac{(I_{Ga+C+O})_{SL} - (I_{C+O})_{Epon}}{(I_{C+O})_{Epon}}\right] \cdot \left[\frac{(\sigma_e)_C}{(\sigma_e)_{Ga}}\right] \qquad (1)$$

Where $(I_{Ga+C+O})_{SL}$ is the total HAADF intensity in the region of the $Ga^+$ implantation and $(I_{Ga})_{SL} \approx (I_{Ga+C+O})_{SL} - (I_{C+O})_{Epon}$ is the approximate intensity of the Ga signal in the line profiles across the block face (Fig. 5b and Supplementary Fig. 6), and $(I_{C+O})_{Epon}$ is the signal originating from carbon and oxygen atoms in the Epon/Araldite embedding material.

Thus, a lower limit for the ratio of Ga to C atoms in the surface layer can be obtained by extrapolating dark-field line profiles from deeper in the FIB-SEM block, i.e., from the constant intensity region on the right side of the line profiles in Fig. 5b and Supplementary Fig. 2, by making the approximation $(I_C)_{SL} \approx (I_{C+O})_{Epon}$. We know, however, that the undamaged Epon/Araldite deeper inside the block contains ~25% the number of oxygen atoms as carbon atoms (Supplementary Tables 1 and 2), and we also know this oxygen is likely to be lost by interaction with electrons as well as by Ga ions in the FIB-SEM. In addition, the FIB milling is a continuous process, which results in complete removal of the surface layer as more ions are incident on the specimen block. This process must therefore involve some $Ga^+$-induced sputtering of carbon atoms. These factors indicate that the concentration of Ga relative to C in the surface layer could be significantly higher than given by the approximation in Equation (1) above. If the constant HAADF STEM signal deeper inside the block represents C and O atoms and oxygen atoms in the surface layer are lost due to radiolysis by the 1.5 keV electron beam, we can write:

$$(I_C)_{SL} = (I_{C+O})_{Epon}\left[1 + \frac{(n_O)_{Epon}(\sigma_e)_O}{(n_C)_{Epon}(\sigma_e)_C}\right]^{-1} \qquad (2)$$

Then, using the composition of Epon in Supplementary Table 1, and the elastic cross sections in Supplementary Table 3, we find that $(I_C)_{SL} = 0.74 \times (I_{C+O})_{Epon}$. This increases the estimated atomic ratio of Ga to C by a factor of 1.36 relative to the value obtained by ignoring loss of oxygen.

**Supplementary Note 2: Estimated composition of the FIB-milled surface layer of the Epon block, based on STEM-EELS imaging of sections cut perpendicular to the block face.**

The atomic ratio of Ga:C in the surface layer (*SL*) of the block $\left(\frac{N_{Ga}}{N_C}\right)_{SL}$ is proportional to the ratio of integrated core-edge signals in energy loss window Δ = 100 eV, above the respective core edges, $\left(\frac{I_{Ga}(\Delta,\alpha)}{I_C(\Delta,\alpha)}\right)_{SL}$ after inverse power law background subtraction, and $\alpha$ = 10 mrad is the EELS collection semi-angle. The ratio of core signals is then multiplied by the inverse ratio of the core-edge cross sections for the same energy range Δ and collection semi-angle $\alpha$, $\left(\frac{\sigma_C(\Delta,\alpha)}{\sigma_{Ga}(\Delta,\alpha)}\right)$ to obtain the atomic ratios, in the surface layer:

$$\left(\frac{N_{Ga}}{N_C}\right)_{SL} = \left(\frac{I_{Ga}(\Delta,\alpha)}{I_C(\Delta,\alpha)}\right)_{SL} \cdot \left(\frac{\sigma_C(\Delta,\alpha)}{\sigma_{Ga}(\Delta,\alpha)}\right) \quad (1)$$

Due to the large energy loss difference between the C K edge (285 eV) and the Ga $L_{2,3}$ edge (1,114 eV), and uncertainly in the Ga cross section near the threshold energy, as well as some uncertainty in the EELS collection semi-angle α, we make use of a standard reference spectrum from gallium oxide ($Ga_2O_3$) that we have acquired. We can then write:

$$\left(\frac{N_{Ga}}{N_O}\right)_{Ga_2O_3} = \left(\frac{I_{Ga}(\Delta,\alpha)}{I_O(\Delta,\alpha)}\right)_{Ga_2O_3} \cdot \left(\frac{\sigma_O(\Delta,\alpha)}{\sigma_{Ga}(\Delta,\alpha)}\right) \quad (2)$$

from which we obtain an experimentally determined estimate for $\sigma_{Ga}(\Delta,\alpha)$ in terms of $\sigma_O(\Delta,\alpha)$:

$$\sigma_{Ga}(\Delta,\alpha) = \sigma_O(\Delta,\alpha). \left(\frac{I_{Ga}(\Delta,\alpha)}{I_O(\Delta,\alpha)}\right)_{Ga_2O_3} \cdot \left(\frac{N_O}{N_{Ga}}\right)_{Ga_2O_3} \quad (3)$$

Then substituting (3) into (1), we obtain the estimated ratio of Ga to C in the surface layer:

$$\left(\frac{N_{Ga}}{N_C}\right)_{SL} = \left(\frac{I_{Ga}(\Delta,\alpha)}{I_C(\Delta,\alpha)}\right)_{SL} \cdot \left(\frac{\sigma_C(\Delta,\alpha)}{\sigma_O(\Delta,\alpha)}\right) \cdot \left(\frac{N_{Ga}}{N_O}\right)_{Ga_2O_3} \cdot \left(\frac{I_O(\Delta,\alpha)}{I_{Ga}(\Delta,\alpha)}\right)_{Ga_2O_3} \quad (4)$$

The ratio of $\sigma_C(\Delta)$: $\sigma_O(\Delta)$ for a primary beam energy of 300 keV and a window of Δ = 100 eV above the C and O K-edges is 3.56:1 from Gatan DM Software with EELS plug-in[34] (Supplementary Figure 1).

The ratio of Ga:O in gallium oxide is 2:3, so that $\left(\frac{\sigma_C(\Delta,\alpha)}{\sigma_O(\Delta,\alpha)}\right) . \left(\frac{N_{Ga}}{N_O}\right)_{Ga_2O_3}$ = 2.39.

For $\Delta$ = 100 eV and $\alpha$ = 10 mrad, the measured value of $\left(\frac{I_O(\Delta,\alpha)}{I_{Ga}(\Delta,\alpha)}\right)_{Ga_2O_3}$ was 3.57 (±0.17) for five measurements (± standard deviation)

Therefore:

$$\left(\frac{N_{Ga}}{N_C}\right)_{SL} = \left(\frac{I_{Ga}(\Delta,\alpha)}{I_C(\Delta,\alpha)}\right)_{SL} \times 8.5$$

#### Supplementary Movies

**Supplementary Movie 1 | 3D image of presynaptic terminal obtained from a sample of cultured neurons in the FIB-SEM.** 3D data set was acquired in a Zeiss Crossbeam 550 FIB-SEM operating at a primary electron beam energy of 1.5 keV and probe current of 2 nA. The sample was prepared following the UC San Diego NCMIR protocol, as described by Deerinck et al.[26]. The sample was milled with a 30 keV $Ga^+$ beam, current of 700 pA, pixel dwell time of 16 µs and a pixel size of 3 nm in x, y and z. A cluster of synaptic vesicles in evident in the animation, each with a hollow lumen and outer diameter of ~40 nm. The data illustrate that features are discerned on a scale of 5-10 nm, which would not be distinguishable without the high surface layer concentration of implanted Ga atoms, and the ability to acquire electron images at an electron fluence of ~$10^4$ electrons/$nm^2$.
